# Fragmentation of extracellular ribosomes and tRNAs shapes the extracellular RNAome

**DOI:** 10.1101/2020.01.29.923714

**Authors:** Juan Pablo Tosar, Mercedes Segovia, Fabiana Gámbaro, Yasutoshi Akiyama, Pablo Fagúndez, Bruno Costa, Tania Possi, Marcelo Hill, Pavel Ivanov, Alfonso Cayota

## Abstract

A major proportion of extracellular RNAs (exRNAs) do not co-isolate with extracellular vesicles (EVs) and remain in ultracentrifugation supernatants of cell-conditioned medium or mammalian blood serum. However, little is known about exRNAs beyond EVs. We have previously shown that the composition of the nonvesicular exRNA fraction is highly biased toward specific tRNA-derived fragments capable of forming RNase-protecting dimers. To solve the problem of stability in exRNA analysis, we developed RI-SEC-seq: a method based on sequencing the size exclusion chromatography (SEC) fractions of nonvesicular extracellular samples treated with RNase inhibitors (RI). This method revealed dramatic compositional changes in exRNA population when enzymatic RNA degradation was inhibited. We demonstrated the presence of ribosomes and full-length tRNAs in cell-conditioned medium of a variety of mammalian cell lines. Their fragmentation generates some small RNAs that are highly resistant to degradation. The extracellular biogenesis of some of the most abundant exRNAs demonstrates that extracellular abundance is not a reliable input to estimate RNA secretion rates. Finally, we showed that chromatographic fractions containing extracellular ribosomes can be sensed by dendritic cells. Extracellular ribosomes and/or tRNAs could therefore be decoded as damage-associated molecular patterns.

## INTRODUCTION

Extracellular RNA (exRNA) profiling in biofluids such as urine, plasma or serum is a promising approach for early disease detection and monitoring in minimally invasive liquid biopsies ^1^. Although plasma cell-free DNA analysis has proven powerful to detect cancer-associated mutations ^2^ or altered DNA methylation events ^3^, exRNA analysis has the potential to inform about transcript expression, post-transcriptional modifications and splicing variants ^4^. Additionally, cells use exRNAs to communicate and reciprocally regulate their gene expression, even in other tissues ^5–7^.

Because RNA is less stable than DNA in biofluids such as plasma due to the high RNase activity of these samples ^8^, exRNAs are typically studied in the context of lipid membrane-containing extracellular vesicles (EVs) ^7,9,10^ or lipoprotein particles (LPPs) ^11,12^. Alternatively, exRNAs can achieve high extracellular stability by their association with proteins^13,14^. However, extracellular soluble ribonucleoproteins remain the least studied exRNA carriers ^15^, with most attention thus far placed at the level of EVs. A new extracellular nanoparticle was recently discovered ^16,17^, but the exact identity and RNA content of these complexes termed “exomers” needs further characterization.

The well documented involvement of EVs ^18^, and EV-encapsulated RNAs ^5,6,19^ in intercellular communication explains part of the bias toward these exRNA carriers. Extracellular vesicles are actively released by living cells by mechanisms which can be experimentally manipulated ^20,21^ and contain surface proteins that can confer tropism for specific cell types or tissues ^22^. Moreover, nucleic acid loaded ^23^ or unmodified ^24^ EVs have shown therapeutic potential and are moving toward clinical use. On the contrary, the origin of extracellular ribonucleoproteins is controversial ^25^ and it is still unclear whether they are released actively from cells ^26^ or they constitute stable intracellular complexes passively released by distressed, damaged or dying cells ^14^.

It is widely accepted that most of the cell-free DNA found in human blood plasma is originated from apoptotic hematopoietic cells ^27,28^. However, the contribution of dead cells to□exRNA□profiles is unclear. Although identification of RNAs actively and selectively released to the extracellular space can help to understand physiological communication circuits between nonadjacent cells, nonselective RNA release (derived from either live or dead cells) can provide cell- or tissue-specific extracellular signatures which can be identified, at least in theory, by deconvolution analysis ^29^. However, it is reasonable to expect that those exRNAs that are not contained inside EVs would be rapidly degraded by extracellular RNases. This is probably the main reason why nonvesicular exRNAs have not attracted much attention until very recently ^16,26^.

Strikingly, a major proportion of extracellular small RNAs are found outside EVs ^13,30^. Furthermore, nonvesicular exRNA profiles are highly biased toward glycine and glutamic acid 5’ tRNA halves. This has been extensively documented both in cell culture media ^30,31^ and in biofluids such as urine, blood serum, saliva or cerebrospinal fluid ^32–35^. The abundance of these species in the extracellular, nonvesicular fraction ^30,34,35^challenges the widespread belief that exRNAs are unstable when not present inside EVs and raise the question on the origin of their remarkable extracellular stability. A possible answer comes from our recent report that glycine and glutamic acid□5’ tRNA halves can form homo- or heterodimeric hybrids which render these structures resistant to single-stranded RNases ^36^. The RNAs with predicted dimer-forming capacity are slightly shorter (30 – 31 nt) than the 5’ tRNA halves generated by endonucleolytic cleavage of the anticodon loop during the stress response (typically 34 – 35 nt) ^37–39^. Interestingly, nonvesicular extracellular fractions are usually enriched in glycine and glutamic acid tRNA halves of precisely 30 – 31 nt ^30,32^, suggesting a causal link between extracellular stability and abundance ^32^.

We hypothesized that extracellular RNA degradation is a major force shifting what cells release to the extracellular space toward those species with higher extracellular stability. Consequently, we reasoned that conventional exRNA profiling fails to capture the complete set of RNAs released from cells to the extracellular space, frustrating attempts to infer RNA secretion mechanisms from comparisons between intracellular and extracellular RNA profiles. To study this, we compared exRNA profiles in cell-conditioned media obtained with or without addition of recombinant ribonuclease inhibitor (RI). Surprisingly, addition of RI greatly increased the complexity of exRNA profiles, providing evidence for the presence of extracellular ribosomes and tRNAs and for their rapid decay to rRNA- and tRNA-derived fragments. Some of these fragments are highly stable and can accumulate in RNase-rich samples such as serum, even when present outside EVs. Overall, we provide a dynamic and comprehensive characterization of the nonvesicular RNAome which can impact biomarker discovery in biofluids. Because our data is consistent with a significant fraction of nonvesicular RNAs being released from distressed, damaged or dying cells, we also provide evidence supporting an immunomodulatory role for some of these extracellular ribonucleoprotein particles.

## METHODS

### Reagents

A full list of reagents including antibodies, primers and probes used in this study are reported in *SI Materials and Methods*.

### Preparation of cell-conditioned medium

Cells were cultured in DMEM + 10% fetal bovine serum (FBS; Gibco) at 37°C and 5% CO_2_. Conditioned medium was typically derived from one 75 cm^2^ flask for chromatographic analysis or from one 150 cm^2^ flask for electrophoresis. Cells were plated at a density which was adjusted to obtain a confluency of 80% at the endpoint of the experiment. Recombinant ribonuclease inhibitor, murine (RI; New England Biolabs) was added in selected experiments at a final concentration of 4-8U / mL, either at the time of medium renewal or following collection of cell-conditioned media.

Preparation of serum-containing conditioned medium (adherent cells; protocol 1): cells were grown in DMEM + 10% FBS (“S+” medium). They were washed with S+ medium, and incubated in S+ for variable periods of time which ranged from 1 to 48 hours.

Preparation of serum-free conditioned medium (adherent cells; protocol 2): cells were plated on day zero and grown in S+ for 24 hours. Later, they were washed with PBS, grown in Mammary Epithelial Growth Medium without antibiotics and without bovine pitutary extract (MEGM, Lonza) for 48 hours, washed again with PBS, and grown in MEGM for additional 48 hours.

Preparation of serum-free conditioned medium (adherent cells; protocol 3): cells were grown in S+, washed with DMEM and incubated with DMEM + 1x Insulin-Transferrin-Selenium solution (“ITS” medium) for one hour.

ExRNA analysis after short washes in PBS or Hank’s Balanced Salt Solution (HBSS) (adherent cells; protocol 4): cells were grown in S+ medium until 80% confluent, washed three times with warm buffer (5 seconds per wash), and washed a forth time for 30 seconds with 5 mL (in T75 flasks) or 10 mL (in T150 flasks) of warm buffer plus 20 – 40 U RI (respectively). Buffers could correspond to PBS, PBS plus divalent cations (PBS+), HBSS or HBSS plus divalent cations (HBSS+) depending on the experiment. Flasks were gently tilted from side to side during washes.

For protocols 1-4: conditioned media or conditioned-buffers were centrifuged at 800 x g at room temperature to remove detached cells and then spinned twice at 4°C and 2,000 x g. The supernatants were either processed immediately or stored at −20°C for later use. If frozen, media was spinned again at 4°C and 2,000 x g upon thaw.

ExRNA analysis after short washes in PBS (suspension cells; protocol 5): cells were grown in S+ medium until the desired cell density, spinned down at 300 x g for 5 min at room temperature, resuspended in DMEM and immediately spinned down again. The cell pellet was then resuspended in PBS (+RI) and immediately spinned down at 300 x g for 5 min. The supernatant (“cell-conditioned PBS”) was centrifuged four times at 2,000 x g and 4°C to remove any remaining cell and used immediately or stored at −20°C.

### Preparation of the nonvesicular extracellular fraction by ultracentrifugation

The cell-condtioned medium already spinned at 2,000 x g was centrifuged for 2.5 h at 100,000 x g and 4°C in a Beckman Coulter Optima XPN-90 ultracentrifuge using a SW40 Ti rotor. The supernatant was concentrated to ∼250 μl with 10,000 MWCO ultrafiltration units (Vivaspin 20, Sartorious Stedim Biotech) and subjected to size-exclusion chromatography or RNA extraction with TRIzol (according to manufacturer’s instructions).

### Size exclusion chromatography

The 100,000 x g supernatants of the cell-conditioned medium were concentrated by ultrafiltration by successive dilutions with PBS to remove phenol red and small molecules. Concentrated non-EV fractions (500 μl) were injected in a Superdex S200 10/300 column (Amersham, GE) and size exlusion chromatography (SEC) was performed at 0.9 ml/min in 0.22 μm-filtered 1x PBS with an Äkta Pure (GE healthcare) FPLC system. All samples were centrifuged at 16,000 × g for 10 min at 4°C before injection in the columns. Fractions of 0.25 mL were collected while monitoring the absorbance at 260 nm and 280 nm. Chromatograms were deconvoluted *in silico* using empirically determined 260/280 ratios from pure bovine serum albumin (BSA) and synthetic tRNA^Gly^ _GCC_ 5’ halves (of 30 nt).

### Sequencing of small RNAs

RNA was purified from selected chromatographic peaks using TRIzol LS and following manufacturer’s instructions. The obtained RNA was diluted in 8 μl of ultra-pure RNase-free water, and 7 μl were used as input for NGS library preparation using the NEBNext Small RNA Library Prep Set for Illumina (New England Biolabs). Sequencing was performed in a MiSeq benchtop sequencer for 200 cycles. The analysis was performed as previously described ^36^ and depicted in *SI Materials and Methods*.

### Proteomic analysis

Fractions corresponding to selected chromatographic peaks were treated with RNase A and later with sequencing grade modified Trypsin (Promega). Peptides were then purified using a C18 ZipTip (Merck Millipore). Eluted peptides were dried in a SpeedVac and resuspended in 10 µL of 0.1 % formic acid. Each sample was injected into a nano-HPLC system (EASY-nLC 1000, Thermo Scientific) fitted with a reverse-phase column (EASY-Spray column, C18, Thermo Scientific). Peptide analysis was carried out in a LTQ Velos nano-ESI linear ion trap instrument (Thermo Scientific). Protein identification was performed with the software PatternLab for Proteomics. Detailed protocols and specificiations are provided in *SI Materials and Methods*. Sample processing and analysis was performed at the Analytical Biochemistry and Proteomic Unit (UByPA) of the Institut Pasteur de Montevideo.

### Northern blotting

RNA samples were run on 10% TBE-urea polyacrylamide gels (ThermoFisher Scientific) and transferred to positively charged nylon membranes (Roche). The membranes were cross-linked by UV irradiation. After cross-linking, the membranes were hybridized overnight at 40°C with digoxigenin (DIG)-labeled DNA probes in DIG Easy Hyb solution (Roche). After low and high stringency washes, the membranes were blocked, probed with alkaline phosphatase-labeled anti-digoxigenin antibody (Roche) and washed with 1x TBS-T. Signals were visualized with CDP-Star ready-to-use (Roche) and detected using ChemiDoc imaging system (BioRad). A detailed protocol and probe sequences are provided in *SI Materials and Methods*.

### Density gradient separations

For separation of ribosomal subunits and ribosomes, a linear gradient composed of 10 – 40 % RNase-free sucrose was used. The gradients were prepared in 20 mM Tris-Cl buffer (pH = 7.4), 4 mM MgCl_2_, 50 mM KCl and 1mM DTT (added fresh). Layers of 40%, 30% 20% and 10% sucrose were added sequentially to a 12 mL polypropylene tube and frozen at −80°C in between. The whole gradient was thawed overnight in the cold-room and used the next day. Extracellular samples were obtained from U2-OS cells using protocol 4 and washing with HBSS+ in the presence of RI. Washes four and five were pooled, concentrated by ultrafiltration and stored at −80°C until use. Concentrated extracellular fractions (0.5 mL) were thawed, layered gently on top of the gradient, and centrifuged at 186,000 x g for 3 hours at 4°C using a SW 40 Ti rotor (acceleration: max; break: min). A density gradient analyzer and fractionator equipped with a Teledyne ISCO UA-6 UV/Vis detector was used to collect fractions of 0.5 mL, starting from the top of the gradient. Collected fractions were treated with 0.5 mL TRIzol to purify both RNA and proteins according to the manufacturer’s instructions.

To separate extracellular vesicles from ribonucleoproteins (RNPs) or other high-density components of the non-EV fraction, high-resolution (12 – 36 %) iodixanol gradients were used following the protocol described in ^26^. Briefly, samples were layered on the bottom of a 12 mL polypropylene tube, and layers of ice-cold 36%, 30%, 24%, 18% and 12% iodixanol (prepared in PBS) were added sequentially on top. The gradients were centrifuged for 15 h at 120,000 x g and 4°C, using a SW40 Ti rotor. Twelve fractions of 1 mL were obtained from the top of the gradient and transferred to new tubes. One half of each fraction was treated with an equal volume of TRIzol and used for RNA and protein purification following manufacturer’s instructions. The other half was twice precipitated with cold (−20°C) 60% acetone, and the pellets were resuspended in 1x loading buffer for Western blot analysis. Loading buffer contained reducing agents or not depending on the primary antibodies intended to use.

### Differentiation of dendritic cells and flow-cytometry

Adherent mouse bone marrow-derived dendritic cells (BMDC) were prepared as described in ^40^. Briefly, bone marrow cells were obtained from the leg bones of C57BL/6 mice and differentiated in culture for eight days in the presence of 0.4 ng / mL GM-CSF. Selected chromatographic fractions or synthetic RNAs were concentrated or diluted to 100 µL (respectively), filter-sterilized, and added to 1 × 10^6^ BMDCs grown in 900 µL of complete medium containing 10% FBS and antibiotics. At t = 24 hs, cells were harvested and analyzed by flow cytometry using a CyAn ADP Analyzer (Dako). A detailed protocol is provided in *SI Materials and Methods*. Levels of IL-1β in the media were measured using a commercial ELISA kit from Biolegend.

## RESULTS

### Addition of RNase A-family inhibitor reshapes nonvesicular exRNA profiles

Chromatographic analysis of the non-EV fraction of MCF-7 cell-conditioned medium (CCM) consistently showed two peaks with Abs 260 > Abs 280 which we termed P1 (V_e_ = 15.0 mL) and P2 (V_e_ = 16.5 mL) (**Figure 1 A; top**). The elution volume of P1 corresponds to that obtained when injecting yeast full-length tRNA^Phe^ or synthetic dsRNAs of 30 nt (**Supplementary Fig. 1, A** and ^36^). In contrast, synthetic ssRNAs of 30 nt elute in the same volume as P2. Treatment of the CCM with RNase A prior to size-exclusion chromatography (SEC) depleted the P1 peak and degradation products were evident with V_e_ ≥ 19 mL (**Figure 1, A; middle**). However, absorbance in the P2 region was only modestly affected. Addition of the recombinant Angiogenin/RNase A-family inhibitor (RI) to the CCM precluded the formation of P2 (**Figure 1, A; bottom**), demonstrating that the P2 peak also contains RNA. In this situation, the P1 peak was still prominent, but quantitatively similar amounts of nucleic acids eluted in the exclusion volume of the column (V_e_ = 7.5 mL; apparent MW ≥ 800 kDa). For simplicity, we will term this new peak “P0”.

**Figure 1:**
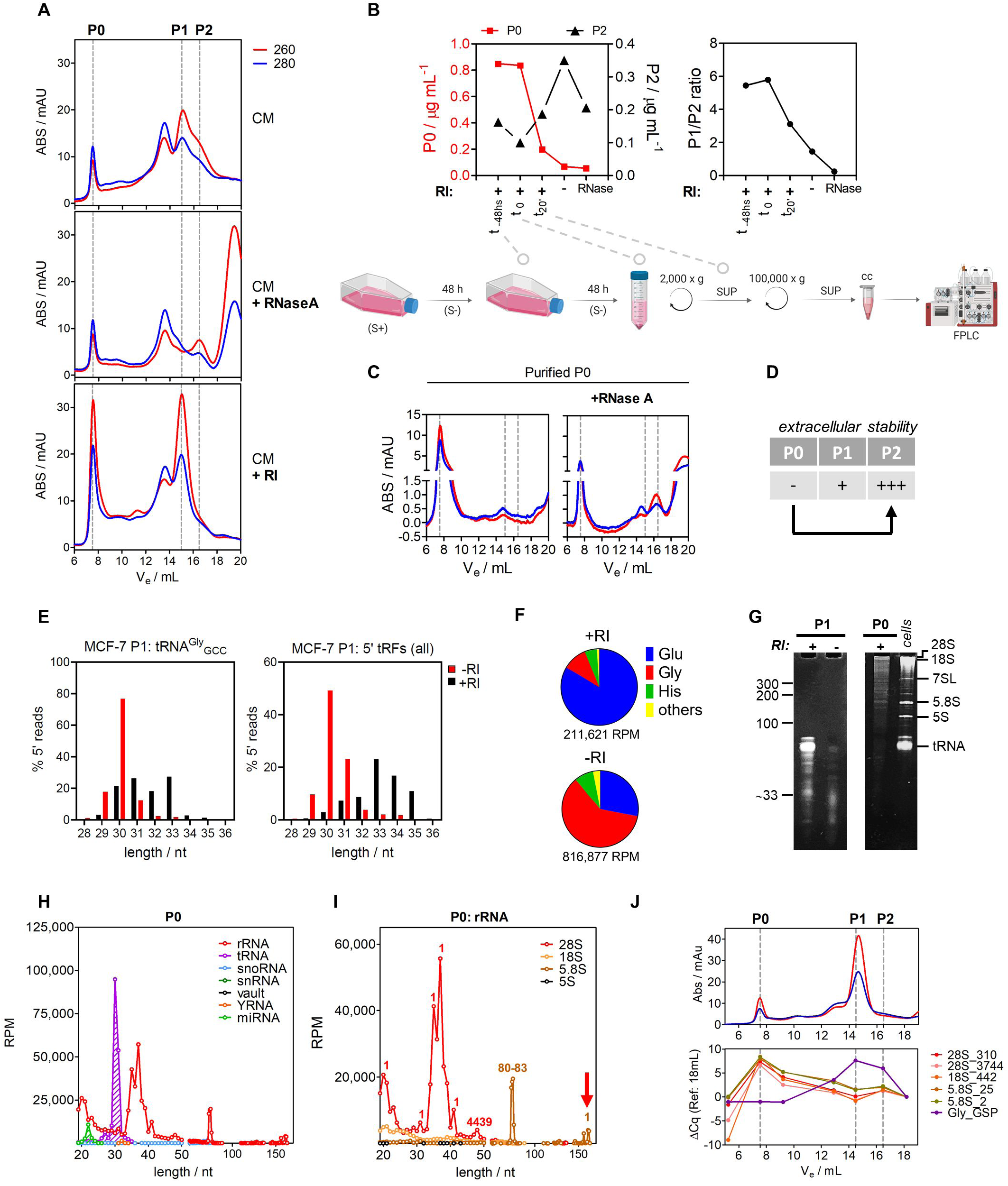
Addition of ribonuclease inhibitor to cell culture conditioned medium (CCM) stabilizes extravesicular ribosomal and transfer RNAs. A) Size exclusion chromatography (SEC) of 100,000 x g supernatants of MCF-7 CCM following addition of RNase A (middle) or Ribonuclease Inhibitor (RI, bottom). Red, Blue: absorbance at 260 nm and 280 nm, respectively. B) The earlier RI was added to the CCM during sample preparation, the higher the P0 peak and the lower the P2 peak (left) and therefore the higher the P1 / P2 ratio (right). C) The P0 peak was purified after SEC and treated with RNase A, which partially reconstituted the P2 peak. D) Comparison of extracellular stabilities of the P0, P1and P2 peaks. E) Size distribution of reads mapping to the 5’ half of glycine tRNA (left) or to all tRNAs (right) in the P1 peak from MCF-7 cells either with (black) or without (red) addition of RI. F) Relative representation of reads mapping to different tRNA isoacceptors in the P1 peak of MCF-7 cells obtained after treatment (top) or without treatment (bottom) of RI. G) Analysis of the P1 peak either with (+) or without (-) RI treatment (left gel) or the P0 peak (right gel) in a denaturing (7M urea) 8% polyacrylamide gel. Sizes were estimated based on a MCF-7 RNA lysate (“cells”) and a RiboRuler Low Range small RNA ladder (left marks; the 33 nt mark was calculated based on Rf). H) Size distribution of small RNA sequencing reads mapping to rRNAs (red), tRNAs (violet) or other ncRNAs (see legend) in the P0 peak of MCF-7 cells. RPM: reads per million mapped reads. I) As in (H), but showing only the reads aligning to rRNAs. The number above each peak denotes the starting position of most reads defining that peak in the corresponding rRNA. J) Amplification by random-primed RT-qPCR of different regions of 28S, 18S and 5.8S rRNAs in different fractions collected after SEC separation of MCF-7 CCM. Gly_GSP: amplification of glycine 5’ halves by using a gene-specific primer during RT. Numbers following rRNA primers (e.g., 28S_310) represent the position of the 5’ end of the expected amplicon.

The high MW RNA-containing complex in P0 was mutually exclusive with the P2 peak (**Figure 1, B**). The earlier the RI was introduced in our purification protocol, the higher the P0/P2 and the P1/P2 ratios. Furthermore, RNase A treatment of the purified P0 fraction generated partial degradation products which eluted in P2 (**Figure 1, C**). Taken together, these results suggest that P2 contains a small RNA population formed by partial fragmentation of high molecular weight RNAs within P0. For some reason, fragments from P2 seem to be highly resistant to the action of RNase A.

The dramatic effects on exRNA profiles induced by early RI addition highlight an often-underappreciated fact in exRNA studies. When differential extracellular stabilities exist (**Figure 1, D**), extracellular abundances are not informative of relative RNA secretion rates. Our results also suggest that MCF-7 cells grown under serum-free conditions mostly release RNA complexes that elute as P0 and P1. Extracellular RNases modify this profile and generate a scenario where P1 and P2 are the main RNA populations, but RNAs in P2 do not seem to be directly released despite being highly abundant in the untreated CCM.

### RI-SEC-seq enables detection of extracellular nonvesicular tRNAs and rRNAs

Small RNA sequencing of P1 obtained with or without addition of RI was previously published by our group ^36^ and showed a predominance of 5’ tRNA halves derived from tRNA^Gly^_GCC_ and tRNA^Glu^_CUC_. However, tRNA-derived fragments in the “+RI” library tended to be longer than in “–RI” (33-35 vs 30-31). When RI was present, there was a predominance of 5’ tRNA^Glu^, and when RI was not present the predominance shifted toward 5’ tRNA^Gly^ (**Figure 1, E-F and Supplementary Fig. 1, B**). These same species were the most abundant small RNAs in the total nonvesicular fraction of MCF-7 CCM ^30^ and are frequently detected in human biofluids ^32^. The ability of these fragments to form RNase-resistant homo and heterodimers ^36^ is consistent with their detection in the absence of RI and their elution in the P1 peak. However, addition of RI recovered a higher diversity of small RNAs in the P1 peak, including fragments of rRNA and full-length snoRNAs. Interestingly, the most frequently detected 28S rRNA fragment corresponds to the region which is hybridized by the most frequently detected snoRNA during ribosomal RNA maturation (**Supplementary Fig. 1, C-D**). This strongly suggests the presence of additional extracellular RNA hybrids, albeit with lower extracellular stabilities than dimers of 5’ tRNA halves.

Sequencing of full-length mature tRNAs is challenging due to the abundance of modified ribonucleotides and their strong self-complementarity. Therefore, specific protocols for tRNA sequencing and analysis have been developed in recent years ^41–44^. Despite using conventional ligation-based library preparation methods, we obtained a low but nonnegligible number of reads corresponding to two full-length tRNA^Glu^ isoacceptors (**Supplementary Fig. 1, E**). These sequences were exclusively detected in the “P1 + RI” library in an almost complete form except from the initial 12 nucleotides and contained the 3’ terminal nontemplated CCA addition indicative of mature tRNAs. Analysis of the P1 + RI fraction in a denaturing RNA gel showed that, indeed, most RNAs migrated like full length tRNAs (**Figure 1, G**). In contrast, the majority of RNAs in “P1 –RI” were < 33 nt. Altogether, these results suggest that exRNA profiles are biased toward the most stable RNAs by the action of extracellular ribonucleases and they do not necessarily reflect the composition of RNAs actually released by cells. Although the P1 peak was detectable with or without RI addition, the RNAs eluting at that volume shifted from a predominance of full-length tRNAs to the highly stable tRNA^Gly^ _GCC_ 5’ tRNA halves of 30 - 31 nt.

The P0 peak contains mostly high-MW RNA, as evidenced by denaturing gel electrophoresis (**Figure 1**, and because reinjection of the purified RNA from the P0 peak still eluted in the void volume (**Figure 1, C**). Small RNA sequencing showed the same tRNA-derived fragments found in P1, but overwhelmingly higher levels of rRNA fragments (**Figure 1H-I**). The majority of these were 28S rRNA 5’ fragments of 20-40 nucleotides. Our size-selected libraries were too short (< 200 nt) to capture the complete 28S (5070 nt) or 18S (1869 nt) rRNAs, although sequences spanning the entire 28S rRNA were detectable (**Supplementary Fig. 1, F**). Nevertheless, the entire 5.8S rRNA could be read (156 nt; 3,108 RPM). Fragments of 72-75 nt corresponding to 5.8S rRNA 3’ halves were also robustly detected (44,991 RPM).

To demonstrate the presence of full-length rRNAs in P0 we performed random-primed reverse transcription and qPCR amplification with primers spanning different regions of 28S, 18S and 5.8S rRNAs (**Figure 1, J**). All primer sets amplified comparably and exclusively in P0, while tRNA^Gly^_GCC_ 5’ tRNA halves peaked in P1 and P2 as expected. This evidence, together with sequencing of the entire 5.8S rRNA, strongly suggests the release of intact rRNAs from both ribosomal subunits to the extracellular space and outside EVs.

### Release of rRNAs and tRNAs occurs promptly after 30 seconds of medium renewal

Cell-conditioned medium contains a variety of RNAs that could be released a few seconds, minutes, hours or days before collection. To clearly differentiate between RNAs directly released from cells and fragments generated by extracellular processing of longer precursors, we combined RI treatment with exRNA profiling shortly after medium renewal. To do this, we performed four subsequent washes of cells with PBS and analyzed the RNA content in the cell-conditioned PBS, with a conditioning time as short as 30 seconds (**Figure 2, A**). Strikingly, both the rRNA-associated P0 and the tRNA-associated P1 peaks were detected in the four washes with very little variation between them (**Figure 2, B)**. Similar results were obtained when washing with serum-free chemically defined growth medium (MEGM; **Supplementary Fig. 2, A)**, albeit with a higher wash-to-wash variability. Thus, detection of rRNAs and tRNAs in the 4^th^ PBS or MEGM wash was not a consequence of carry-over from previously longer incubations. Instead, fast release of these RNAs occurred every time the cells were washed. Increasing incubation time from 30 seconds to 10 minutes did not increase extracellular RNA levels (**Figure 2, C-D**).

**Figure 2:**
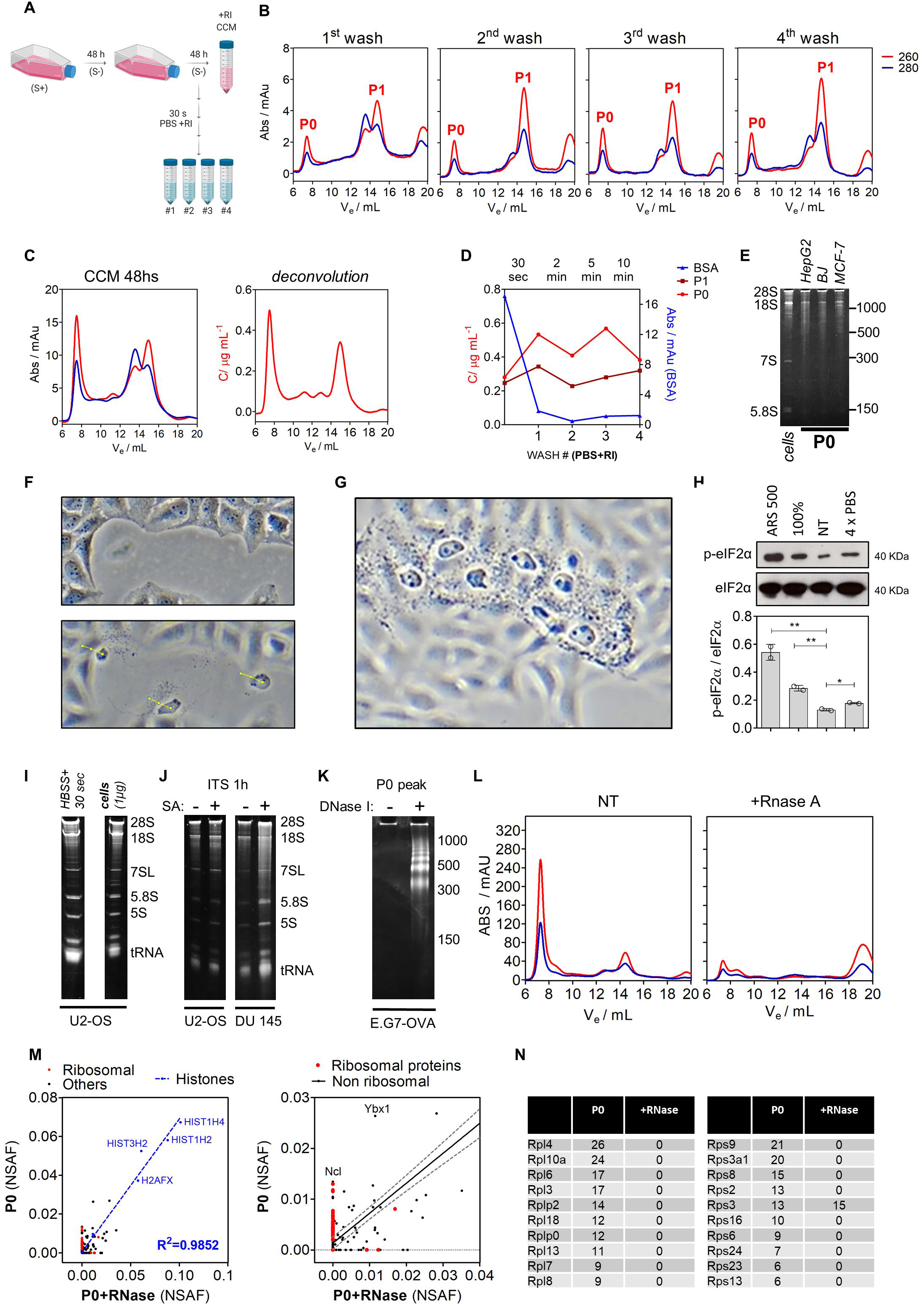
Intracellular nucleic acids are released to the extracellular space after short washes in serum-free media or isotonic buffers. A) Schematic representation of the experimental protocol used in panels B-E. S(+): DMEM + 10% FBS. S(-): MEGM media, serum-free. B) RNA analysis by SEC in PBS washes #1 to #4 of MCF-7 cells. Conditions identical to those used in Figure 1. C) An example of deconvolution analysis (Abs 260/280 ratio-to-RNA-concentration conversion) applied to a representative chromatogram of MCF-7 CCM (ultracentrifugation supernatant). D) Variation of RNA concentration corresponding to the P0 (light red) and P1 (dark red) peaks in PBS washes #1 to #4. The variation in the Abs 280 nm at the BSA peak is plotted in the right Y axis. Initial values correspond to those present in the CCM. E) Denaturing electrophoresis (7M Urea, 6% PAGE) of the concentrated P0 peak from the 4^th^ PBS wash of HepG2, BJ and MCF-7 cell lines. No RNA purification was performed. “Cells”: MCF-7 RNA lysate. F) U2-OS cells after incubation in serum-free DMEM plus ITS supplement for one hour. Top: monolayer. Bottom: tracking of floating nuclei (yellow line) in three consecutive shots taken one second apart from each other. G) Cluster of floating nuclei. Same conditions as in (F). H) Analysis of eIF2 alpha phosphorylation (Wblot) in nontreated (NT) MCF-7 cells, in cells exposed to four consecutive PBS washes (4 x PBS), in cells cultured after confluency (100%) or exposed to 500 µM sodium arsenite for one hour (ARS 500). Bottom: densitometry analysis in two independent biological replicates of the experiment. I) Denaturing electrophoresis (10% PAGE) of TRIzol-purified total extracellular RNA from U2-OS cells washed for 30 seconds with HBSS, ran alongside 1 µg of purified intracellular RNA from the same cell line. J) Denaturing 10% PAGE exRNA analysis in U2-OS (left) or DU 145 (right) cell-conditioned media (ITS, one hour) obtained in the presence (+) or absence (-) of 200 µM sodium arsenite. K) Denaturing electrophoresis of the purified P0 peak from E.G7-OVA cells, either treated (+) or not treated (-) with recombinant RNase-free DNase I. Sample preparation after SEC was the same as in panel (E). L) Chromatograms of cell-conditioned PBS form E.G7-OVA cells. The sample was separated into two aliquots, one of which was treated with RNase A before SEC (right). M) Proteomic analysis of the RNase-treated or not-treated (NT) P0 peak from E.G7-OVA cells. Blue: histones. Red: ribosomal proteins. NSAF: normalized spectral abundance factor. N) List of the top ten proteins from the large (left) and small (right) ribosomal subunits producing the higher number of spectra in the proteomic analysis of the P0 peak from E.G7-OVA cells.

To study whether this phenomenon is generalizable, we repeated the same experiments in human malignant cell lines derived from a variety of tissues (MCF-7, U2-OS, HepG2), in normal human dermal fibroblasts cultured at low passage (BJ), and in cell lines derived from mice (LL/2) or nonhuman primates (Vero). Even though cell-normalized RNA quantities varied in each case, the presence of the P0 and P1 peaks in the 4^th^ PBS was h was highly reproducible (**Figure 2, E and Supplementary Fig. 2, B)**.

Quantitatively, RNA levels in each wash were 0.1 – 0.5 % of the total RNA present in the culture (estimated by multiplying cell number by average RNA contents per cell). Mechanistically, this can be achieved by either 0.5% of dead cells releasing 100% of their cytoplasmic content, or 100% of cells secreting 0.5% of their transcriptome. In this respect, it is tempting to speculate that the presence of high MW ribonucleoproteins in the extracellular medium of cell lines should be the consequence of damaged or dying cells, as some of these cells were observed even after short washes with serum-free media or buffers (**Figure 2, F-G**). As expected, the vast majority of cells were not affected by the washing process and retained their anchorage (**Supplementary Fig. 2, C**). Intracellular levels of phospho-eIF2α (**Figure 2,H**, were only slightly higher than in nontreated cells (fold change = 1.3 vs. 4.1 in the positive control), even after washing cells with PBS. Because eIF2α phosphorylation is a hub of various stress response programs ^45^, a rapid reprogramming of intracellular RNA levels is not expected as a considerable outcome of our washing protocol, which does not induce stress. Furthermore, cold (4°C) HBSS+ washes did not impaired exRNA detection (data not shown) arguing against an active ATP-dependent secretion mechanism.

### Extracellular nonEV RNAs mirror the intracellular transcriptome

To distinguish between selective and nonselective release of cytoplasmic contents, we scaled up our cultures in an attempt to detect less abundant nonvesicular exRNAs. Surprisingly, denaturing PAGE analysis of TRIzol-purified RNA from intracellular and extracellular fractions showed virtually identical results in U2-OS cells (**Figure 2, I**) and other adherent cell lines. These include, among others, the DU145 cell line which is deficient in ATG5 and therefore in autophagy ^46^, but still showed extracellular rRNAs, 7SL RNAs and tRNAs when incubated in serum-free media supplemented with growth factors for just one hour (**Figure 2, J**). As a consequence, the release mechanism operating herein seems to be unrelated to the autophagy-dependent (presumably ATG5-mediated) secretion of cytoplasmic nucleic acids described in ^26^. All major intracellular RNA classes were detectable in the extracellular space and the relative abundance between classes was also conserved. Selectivity within a given RNA class (e.g., tRNAs) will be studied in in later sections of this work.

In order to extend our analysis to cells in suspension, slight modifications in our washing protocol were introduced. Despite these modifications, the chromatograms of the RI-treated cell-conditioned PBS from the human THP-1 (**Supplementary Fig. 3, A**) and the murine EG.7-OVA cell lines showed the characteristic P0 and P1 peaks. The washing protocol itself did not induce apoptosis. On the contrary, there was a significant reduction in the percentage of Annexin V-positive, propidium iodide-positive late apoptotic cells recovered after the washes (**Supplementary Fig. 3, B-C**), suggesting that not all apoptotic cells could be collected by low speed centrifugation and at least some cells released their contents into the medium. Consistent with this, and in contrast to what was previously observed in adherent cell lines, the P0 peak clearly contained fragmented DNA (**Figure 2, K**). Proteomic analysis showed that histones were the most abundant identifiable proteins in the P0 peak, and their association with this peak was not affected by RNase A treatment of the sample before SEC (**Figure 2, L-M**). In contrast, ribosomal proteins were almost completely depleted from the P0 peak when the medium was treated with RNase A (**Figure 2, M-N**). Detection of most ribosomal proteins in a high molecular weight complex with an RNA scaffold strongly suggests the elution of ribosomes or ribosomal subunits in the P0 peak from these suspension cells.

In summary, even after washing cells to remove RNAs released to the medium after prolonged incubations, the most abundant intracellular nucleic acids and their associated proteins could be detected in the extracellular space. For cells in suspension, the fragmentation pattern of exDNA suggests that apoptotic cells are a main source of extracellular nucleic acids, despite these cells being lowly represented in the culture. In adherent cells, the apparent lack of detectable exDNA could be explained by the induction of a form of cell damage which preserves nuclear membrane integrity in a limited number of cells exposed to serum-free media (**Figure 2, F-G**).

### Identification of ribosomes and oligoribosomes in the extracellular space

Proteomic analysis of the P0 peak from EG.7-OVA cells strongly suggested the presence of both nucleosomes and ribosomes (**Figure 2, K-N**) which were presumably derived from apoptotic cells. Because DNA was absent from the P0 peak of adherent cell lines exposed to a series of brief washes with buffers or serum-free media, the mechanisms responsible for RNA release in adherent and suspension cells seemed to be essentially different. However, we could identify features with size and morphology reminiscent of ribosomes by negative-stain transmission electron microscopy of the concentrated P0 peak purified from adherent Hep G2 cells (**Supplementary Fig. 3, D**).

To confirm the presence of extracellular ribosomes in other adherent cell lines, we resort to study concentrated extracellular fractions by velocity sedimentation in sucrose gradients optimized for polyosome preparations (**Figure 3, A**). Analysis of the velocity gradients showed three clearly defined 254 nm peaks in the region corresponding to ribosomal subunits or ribosomes (**Figure 3, B**). These peaks corresponded to the small ribosomal subunit 40S (low levels of 28S, 5.8S and 5S rRNA, high levels of 18S rRNA, low levels of RPL7a, high levels of RPS6), 80S monosomes (all rRNAs and all ribosomal proteins detectable with the expected stoichiometry), and, considering a lack of detectable ribosomal proteins in fraction #10 and their reappearance in fraction #11, two ribosomes (2x). Interestingly, a small 254 nm peak in fraction #14 was accompanied by a faint but detectable band for RPL7a and RPS6 and was indicative of oligoribosomes or polysomes. These were further stabilized by treating cells with the translation elongation blocker cycloheximide (but not with the premature terminator puromycin) straight before HBSS+ washes (**Figure 3, C**).

**Figure 3:**
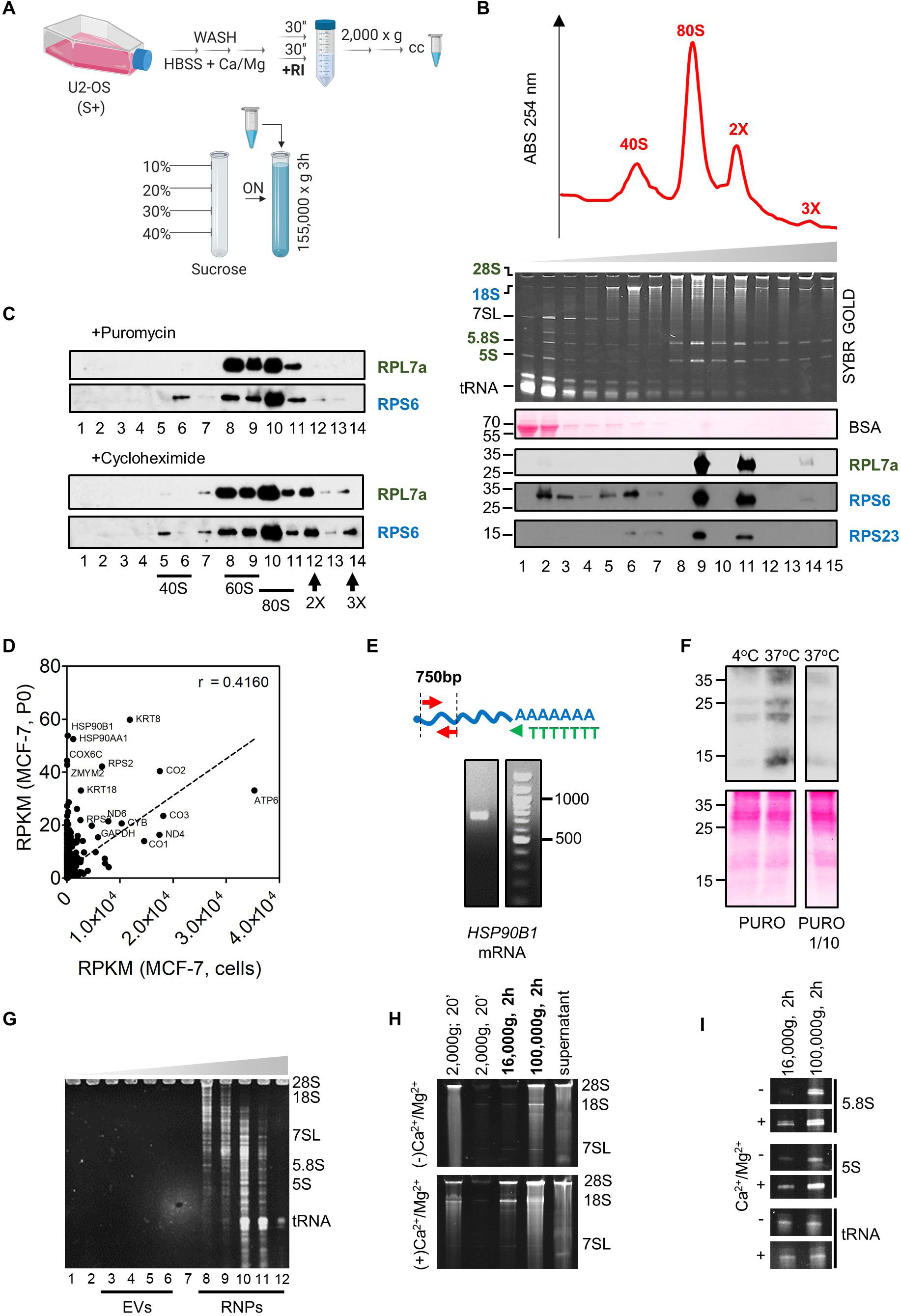
The extracellular space contains ribosomes. A) Schematic representation of the experimental protocol used in panels B-C. B) Velocity gradient (10 – 40 % sucrose, 3 hours at 155,000 x g) of cell-conditioned HBSS (plus divalent cations and RI) from U2-OS cells. Fractions of 0.5 mL were taken out from the top of the tube and treated with TRIzol. RNA was analyzed in a denaturing 10% PAGE and proteins were analyzed by Western blot using antibodies specific to human ribosomal proteins. C) Same as (B), but cells were treated with puromycin (2 µg / mL; 10 min) or cycloheximide (10 µg / mL; 30 min) before washes. D) Analysis of mRNAs in our sequencing of small RNAs from the P0 peak of MCF-7 cells (related to Figure 1H). The mRNA abundances (expressed as reads per kilobase of transcript, per million mapped reads, RPKM) were estimated from sequenced fragments assuming they represent random fragmentation of their parental mRNAs. Intracellular abundances in MCF-7 cells (“cells”) were based on the Human Protein Atlas (proteinatlas.org) transcriptomic data. E) Amplification by RT-PCR of human HSP90B1 mRNA with an oligo (dT)_18_ RT primer and PCR primers complementary to exons 5 and 9 (2229 bp from 3’ end). F) U2-OS cell-conditioned HBSS+ was treated with 5 µg / mL (left) or 0.5 µg / mL puromycin for two hours at either 37°C or 4°C in the presence of ATP/GTP. Extracted proteins were analyzed by Western blot with an anti-puromycin antibody. G) Isopycnic centrifugation in a 12 – 36 % iodixanol gradient of the concentrated 4^th^ wash of U2-OS cells for 30 seconds with HBSS+. Twelve fractions of 1 mL were collected from the top of the tube and RNA was purified from each fraction and run in a 10% denaturing gel. H-I) U2-OS cells were washed with HBSS containing or not containing divalent ions for 30 seconds. The cell-conditioned HBSS was centrifuged twice at 2,000 x g, followed by sequential ultracentrifugations. Pellets were resuspended in water and, together with the concentrated 100,000 x g supernatant, ethanol precipitated, resuspended, and analyzed by denaturing electrophoresis. Panels H and I show different regions of the same gel.

Detection of 80S particles in puromycin-treated cells can be explained by the tendency of ribosomal subunits to re-associate *in vitro* in the presence of tRNAs ^47^. However, this cannot explain higher order aggregates which were detected deeper in the gradient. Because polysomes are formed by ribosomes sitting on messenger RNAs, we wondered whether the extracellular fractions also contained mRNAs. A reanalysis of our small RNA sequencing in the P0 peak from MCF-7 cells revealed a variety of reads unambiguously mapping to coding sequences. Once again, their extracellular representation was strongly correlated (r = 0.416, p < 0.0001) to their expected intracellular levels, and included mRNAs transcribed from the mitochondrial genome (**Figure 3, D**). To demonstrate the presence of complete, nondegraded mRNAs, RT-PCR was done using the P0 peak as input material and an oligo dT reverse transcription primer. The PCR primers were designed to amplify a 750 bp region at the 5’ end of the HSP90B1 mRNA, one of the most abundantly detected mRNAs in our sequencing study. Bands of the expected size were obtained in MCF-7 and BJ cells (**Figure 3, E** and data not shown).

Next, we addressed whether extracellular ribosomes/polysomes were functional. To do this, we concentrated HBSS+ washes and added ATP/GTP plus puromycin (5 µg / mL) and incubated the samples for two hours at either 4°C or 37°C (**Figure 3, F**). Surprisingly, Western blot analysis with an anti-puromycin antibody showed significant incorporation of the antibiotic to nascent peptides when samples were incubated at 37°C but not at 4°C. To the best of our knowledge, this is the first evidence of the peptidyl transference reaction occurring in extracellular samples obtained without deliberated cell lysis steps.

### Nonvesicular ribosomal RNAs co-purify with extracellular vesicles

Isopycnic centrifugation of exRNAs obtained after washing U2-OS cells with HBSS+ for only 30 seconds confirmed that virtually all RNAs detectable by SYBR gold staining were not associated with extracellular vesicles (**Figure 3, G and Supplementary Fig. 3, E**). However, sequential ultracentrifugation of cell conditioned HBSS (either with or without divalent cations) showed sedimentation of rRNA- and tRNA-containing complexes at speeds usually used to purify small EVs (100,000 x g; **Figure 3, H**). Strikingly, presence of divalent ions favored rRNA sedimentation at speeds typically used to collect large EVs (16,000 x g). This effect was not observed for tRNAs (**Figure 3, I**).

The effect of divalent cations on ribosomal RNAs was also evidenced by SEC analysis. Washing of cells with buffers containing calcium or magnesium depleted the P0 while not affecting the P1 peak (**Supplementary Fig. 3, F-H**). This effect was reproduced when cells were washed in the absence of divalent cations, but these were added immediately afterwards (data not shown). Thus, divalent cations affect the structural integrity of the RNA complexes present in P0. Because we routinely spin samples at 16,000 x g before SEC and given the ultracentrifugation profile of nonvesicular exRNAs (**Figure 3, H-I**), the Ca^2+^/Mg^2+^-induced loss of the P0 peak can be explained by a disassembly of higher order ribosomal aggregates in the absence of divalent cations. This is consistent with the involvement of magnesium ions in the stabilization of ribosomes and polysomes and their disassembly upon EDTA treatment ^48^, although SEC-purification of polysomes is possible under optimized conditions ^49^.

More importantly, our results show that two of the most popular methods for the purification of extracellular vesicles, i.e., SEC and ultracentrifugation ^50^, cannot separate EVs from nondegraded extracellular ribosomes or polysomes.

### Extracellular biogenesis of extracellular tRNA halves

We have previously shown (**Figure 1**) that full-length tRNAs comprise the majority of RNAs present in the P1 peak in RI-treated CCM, but this profile shifted toward RNase-resistant glycine tRNA halves (30-31 nt) in the absence of RI. One possibility is that these tRNA halves were released directly from cells and accumulated because of their resistance to degradation. Alternatively, they could be generated by the fragmentation of extracellular tRNAs.

To distinguish between both possibilities, U2-OS were incubated for one hour in serum-free media (ITS; protocol: 3). The CCM was then split in two aliquots and one of them was treated with RI. Both aliquots were then incubated for 24 hours at 37°C (“cell-free maturation step”; **Figure 4, A**). Isopycnic centrifugation in high-resolution iodixanol gradients confirmed that most of the RNAs were present in the nonvesicular fractions under these experimental conditions (**Figure 4, B**). In both aliquots, extracellular RNAs were similarly and significantly affected by prolonged incubation at 37°C, with a complete loss of 28S and 18S rRNA bands. In contrast, tRNAs were affected to a much lower extent (**Figure 4, C**). Surprisingly, Northern blot analysis showed marked differences between tRNAs. For instance, while tRNA^Lys^ _UUU_ was detectable in both aliquots, tRNA^iMet^ _CAU_ was sensitive to RI addition and tRNA^Gly^ _GCC_ was barely detectable (**Figure 4, D**). The opposite trend was evident for their corresponding 5’ fragments, with tRNA^Gly^ _GCC_ > tRNA^iMet^ _CAU_ > tRNA^Lys^ _UUU_

**Figure 4:**
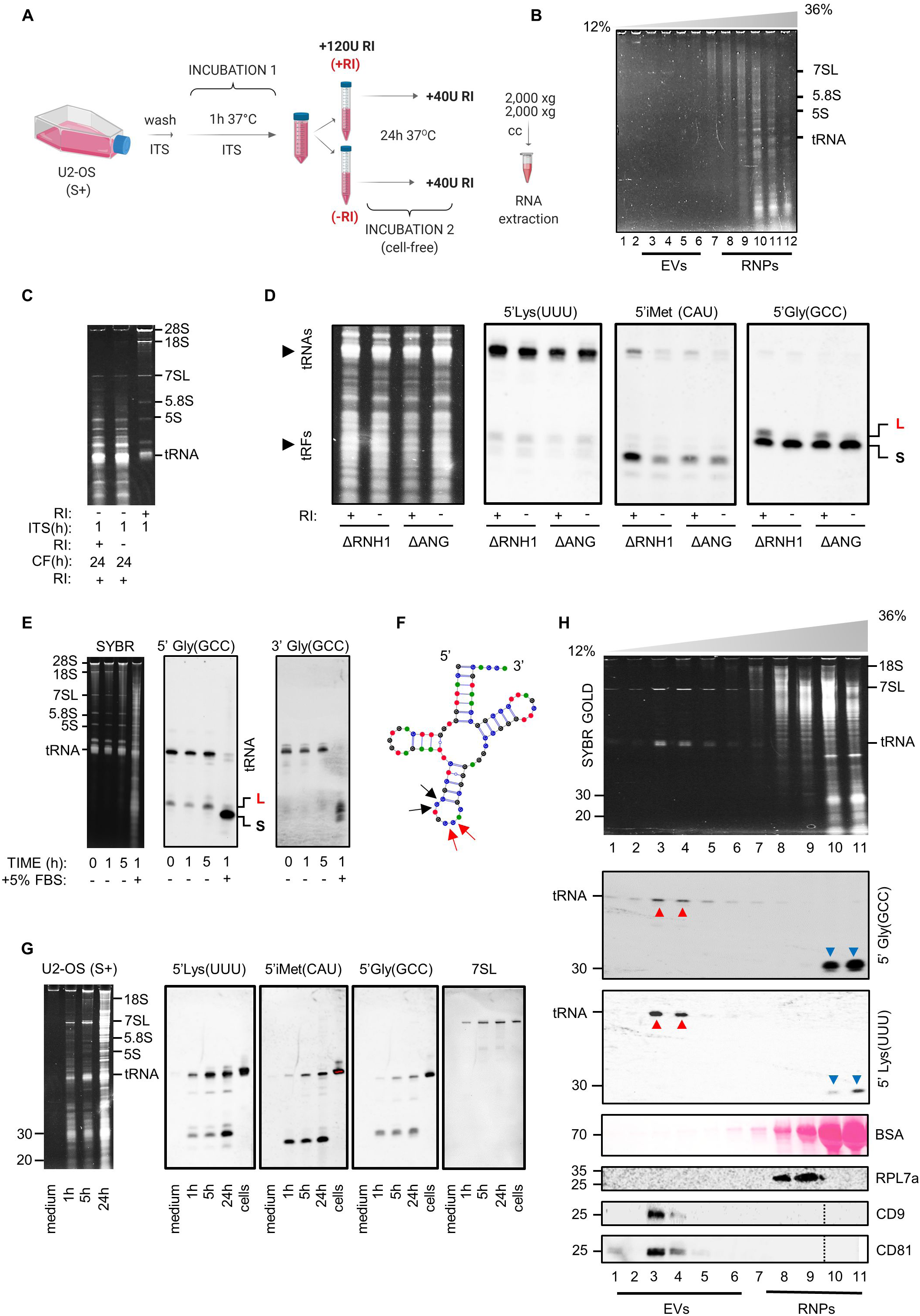
Extracellular tRNAs are processed to extracellular tRNA-derived fragments. A) Schematic representation of the experimental setup used in panels B-D. U2-OS cells were incubated for one hour in ITS medium (protocol: 3) and the CCM was divided into two aliquots, one of which received RI. Both were then incubated at 37°C for 24 hours before exRNA analysis. B) Iodixanol gradient showing most exRNAs were present in the nonvesicular fractions (RNPs) in the input sample (before cell-free processing). C) Analysis of exRNAs by denaturing electrophoresis before and after the cell-free maturation step. D) Northern blot analysis with probes targeting the 5’ region of different tRNAs in samples obtained as explained in (A). U2-OS lacking functional ANG or RNH1 genes were used. L: 5’ tRNA halves of 33 – 34 nt; S: 5’ tRNA halves of 30 – 31 nt. E) Samples were obtained after a short (30 sec) wash of U2-OS cells with HBSS+ (without RI addition), and incubated cell-free at 37°C for 0, 1 or 5 hours, or for 1 hour after addition of S+ medium in a 1:1 ratio. Northern blot was performed with two different probes targeting both halves of tRNA^Gly^_GCC_. F) Cloverleaf representation of glycine tRNA (GCC anticodon, isodecoder #2) with red arrow showing predicted ANG cleavage at the anticodon loop (sequence CpCpA), rendering a 33 or 34 nt 5’ fragment, and black arrows showing a putative alternative cleavage site (sequence CpCpU), generating 30 – 31 nt 5’ fragments. G) Analysis of exRNAs in U2-OS CCM (1, 5 and 24 hours, S+ medium). The concentrated nonconditioned medium was run as a control. Northern blot was done with the same probes as in panel (D), plus a 7SL RNA-specific probe. H) Isopycnic centrifugation of U2-OS CCM (t = 24 hours in S+ medium). Vesicular fractions correspond to those positive for CD9 and CD81 by Western blot and show clear 7SL RNA and full-length tRNA bands which are also confirmed by Northern blot. In contrast, the non-EV fractions are enriched in tRNA-derived fragments.

The 5’ fragments of tRNA^Gly^ were either 33 – 35 nt (i.e., 5’ tRNA halves cleaved at the anticodon loop) or 30 – 31 nt (**Figure 4, D** and **Supplementary Fig. 4, A**). For simplicity, we will call these fragments long tRNA halves (L-tRNAh) and short tRNA halves (S-tRNAh), respectively. Both species are detectable inside U2-OS cells stressed with sodium arsenite but L-tRNAh are produced at much higher levels ^51^. In contrast, only the S-tRNAh are predicted to form RNase-resistant homodimers according to our previous studies ^36^. Strikingly, only the S-tRNAh were detectable in the extracellular milieu in the absence of RI, suggesting that they are indeed very stable. None of these fragments was directly produced by the ribonuclease Angiogenin because nearly identical results were obtained in ΔANG or ΔRNH1 (Angiogenin inhibitor) cells.

Five prime tRNA halves were termed “stress-induced tRNA-derived fragments” or tiRNAs ^38^ because their production is induced by cellular stress. Loading of one microgram of intracellular RNA from U2-OS cells did not show production of these tRNA halves or tiRNAs above the assay’s sensitivity (**Supplementary Fig. 4, A**). If present, intracellular tiRNA levels were lower than the levels of the pre-tRNA^Gly^ _GCC_, as expected in cells not deliberately exposed to stress. Despite the full-length tRNA^Gly^_GCC_ was not detectable after 24 hours of cell-free maturation of the CCM (**Figure 4, D**), the full-length tRNA as well as the L-tRNAh and S-tRNAh were clearly present in the extracellular space if the cell-free maturation step was omitted (**Supplementary Fig. 4, A**). Interestingly, 3’ tRNA-derived fragments were also detectable in the same samples although they are rarely found intracellularly, even in arsenite-treated U2-OS cells ^38^. The higher the degradation state of the exRNA population, the higher the fragment-to-tRNA ratio for both 5’ and 3’ fragments (**Supplementary Fig. 4, B**).

To obtain direct evidence for the conversion of extracellular tRNAs into tRNA halves, we briefly washed U2-OS cells with HBSS+ in the absence of RI and divided the cell-conditioned buffer into four aliquots. These aliquots were incubated for 0, 1 or 5 hours at 37°C before addition of RI and subsequent analysis by Northern blot (**Figure 4, E**). The 4^th^ aliquot was mixed 1:1 with S+ medium (to obtain a final serum concentration of 5%) and incubated for one hour at 37°C. The full-length tRNA^Gly^ _GCC_ and the L-tRNAh (but not the S-tRNAh) were present at t = 0. Incubation for 5 hours at 37°C showed a slight decrease in the intensity of the full-length band and a concomitant increase in L-tRNAh. Strikingly, incubation for only one hour in the presence of serum (RNase-rich sample) entirely converted the full-length tRNA band to S-tRNAh. The L-tRNAh band was also lost.

We wondered whether the L-tRNAh are a necessary intermediate for the formation of extracellular S-tRNAh, or whether these could be formed by an alternative cleavage site at the beginning of the anticodon loop of glycine tRNAs (**Figure 4, F**). *In vitro* controlled digestion with synthetic mimics of 34 nt tRNA^Gly^_GCC_ 5’ halves showed that these RNAs are not preferentially converted to 30-31 nt fragments by bovine RNase A (**Supplementary Fig. 4, C**). This suggests that different tRNA halves might be generated by alternative cleavage sites, as has been recently shown for shorter fragments ^52^.

In summary, evidence from Northern blot (**Figure 4, A-E**) and small RNA sequencing (**Figure 1, E**) consistently show the depletion of full-length tRNAs and L-tRNAh and the concomitant accumulation of glycine S-tRNAh in the presence of serum or in the absence of added RI. Thus, the ubiquitous presence of these specific fragments in biofluids ^32^ can be explained by a combination of factors: a) high expression of tRNA^Gly^ _GCC_ in cells ^42^, b) high susceptibility of tRNA^Gly^ _GCC_ to extracellular nucleases (**Figure 4, D**), c) high resistance of the extracellularly-generated 30 – 31 nt 5’ tRNA halves (S-tRNAh) to degradation ^36^ (**Figure 4, E**) and d) the capacity of these fragments to be sequenced by standard methods (i.e., lack of predictable “hard-stop” modified bases ^41^).

### EVs contain full-length ncRNAs while the non-EV fraction is enriched in ncRNA fragments

All previous assays were performed either in serum-free media or after very short washes with serum-free media or buffers. To evaluate exRNA profiles under standard serum-containing growth conditions (S+; protocol: 1), we collected CCM at different time points, separated vesicles from nonvesicular RNAs by iodixanol gradients and analyzed exRNAs by Northern blot. As depicted in **Figure 4, G**, the S+ medium alone did not reveal detectable bands which could interfere with our analysis.

One striking difference in respect to what was previously observed in ITS incubations (protocol: 3; **Figure 4, B**) or HBSS+ washes (protocol: 4; **Figure 3, G**) was that the tRNA and the 7SL RNA bands were enriched according to what could be expected by rRNA band intensities (**Supplementary Fig. 4, D**). Here, addition of RI (120U; 12U / mL) did not have any observable effects on exRNA profiles.

Having observed that tRNA^Gly^_GCC_ is particularly susceptible to the action of serum RNases (**Figure 4, D-E**), we were surprised to detect it by Northern blot in the extracellular space of U2-OS cells incubated in the presence of 10% serum (**Figure 4, G**). Furthermore, the intensity of the full-length tRNA band was positively correlated to the length of the incubation and there was no interference of FBS-derived tRNAs at the level of sensitivity of the assay. Density gradient analysis reconciled these and previous observations. In serum containing CCM (conditioned for 24 hs) all the observed tRNAs and 7SL RNAs were associated with EVs (**Figure 4, H**). Association of full-length tRNAs and 7SL RNAs with EVs was still observed in ΔYB-1 U2-OS cells (**Supplementary Fig. 4, E**), which is remarkable given the reported involvement of this RNA-binding protein in selecting tRNAs for EV-dependent secretion ^53^. In contrast, neither glycine nor lysine tRNAs were detectable in the nonvesicular fractions. Conversely, tRNA-derived fragments (glycine S-tRNAh in particular) were only present outside EVs.

A similar tendency was found for other RNA polymerase III transcripts like YRNAs and their fragments. Analysis by SL-RT-qPCR showed amplification of full-length YRNAs in 100,000 x g pellets of U2-OS CCM (EV-enriched fraction), but not in the supernatants (EV-depleted fraction). In contrast, YRNA fragments selected from previous extracellular sequencing studies ^30^ as well as miR-21-5p were amplified in both samples at comparable levels (**Supplementary Fig. 4, F**). Similar results have been reported in nematodes, where full-length YRNAs were found exclusively inside EVs whereas their fragments were found outside ^54^.

In summary, long transcripts (including tRNAs) cannot resist prolonged incubation in samples with high RNase activities such as those containing serum except when protected inside EVs. However, some of their fragments are much more resistant and are consistently detected in the non-EV fraction. These fragments define the extracellular nonvesicular RNAome in the absence of added RI. In addition, the low correlation between intracellular and extracellular small RNA profiles can be explained by the fact that many of these fragments are not directly released by cells.

### Immunoregulatory potential of extracellular nonvesicular RNAs

The innate immune system is equipped with a battery of membrane-bound and cytoplasmic pattern recognition receptors (PRRs) capable of sensing pathogen- or damage-associated molecular patterns (PAMPs, DAMPs). Among them, ssRNAs and dsRNAs can be sensed by a variety of PRRs including RIG-1, MDA5, TLR3, TLR7 and TLR8. On the other hand, extracellular exposure of some endogenous intracellular proteins and nucleic acids can also trigger immune cell activation, because exposure of these molecules is interpreted as a sign of cellular or tissue damage ^55^. In this line of reasoning, we wondered whether innate immune cells could sense and react to extracellular ribosomes, tRNAs or their fragments and whether at least some of these RNAs could be considered as DAMPs.

We first observed that the amount of extracellular, nonvesicular RNAs could be increased by the addition of cytotoxic compounds (data not shown). Next, we studied whether dendritic cells (DCs) could sense and react to nonvesicular exRNAs. These cells are regarded as the sentinels of the immune system and link innate and adaptive immune responses ^56^. Thus, we reasoned that if exRNAs are non-silent from an immunological perspective, they should be sensed by DCs in the first place.

ExRNAs obtained from MCF-7 cells were either treated or not with RNase A and later separated by SEC in order to obtain the following samples/fractions: P0, P0 post RNase-treatment and P1. These fractions were concentrated, filtered and added directly to the media of freshly prepared non-primed bone marrow-derived murine dendritic cells (BMDC) (**Figure 5, A-B**). The synthetic dsRNA analogue Poly(I:C) was used as a positive control. After incubation for 24 hours, BMDC maturation was evaluated by flow cytometry monitoring the percentage of CD11c-positive cells expressing high levels of the activation-induced markers MHC class II and CD80 (**Figure 5, C and Supplementary Fig. 5**). The Poly (I:C) present at either 3 or 30 µg / mL in the extracellular space elicited a significant increase in BMDC maturation, compared to nontreated (NT) cells or cells exposed to synthetic single-stranded oligonucleotides (**Figure 5, D**). Interestingly, the purified P0 peak diluted to an RNA concentration as low as 12 ng / mL was sufficient to trigger BMDC maturation. Undiluted P0 (1.2 µg / mL) was highly cytotoxic, with more than 90% of cells staining positive for PI. Strikingly, high levels of the pro-inflammatory cytokine IL-1β were found in the media of these cells (**Figure 5, E**). Altogether, these observations suggest that components in the P0 peak can be sensed by DCs when present in the extracellular space. They can trigger DC maturation and, at higher concentrations, a form of cell death presumably related to over activation or pyroptosis. The latter effect is dependent on RNA because undiluted RNase-treated P0 did not induce IL-1β release nor it triggered significant BMDC maturation in viable cells. These observations strongly argue against potential endotoxin contamination of the P0 fraction that may have led to DC maturation and/or IL-1β secretion. These results afford new avenues in the biological characterization of exRNAs, suggesting at least some of these RNAs are immunologically non-silent. This supports the possibility of an immune surveillance mechanism involving exRNAs or RNP complexes such as extracellular ribosomes.

**Figure 5:**
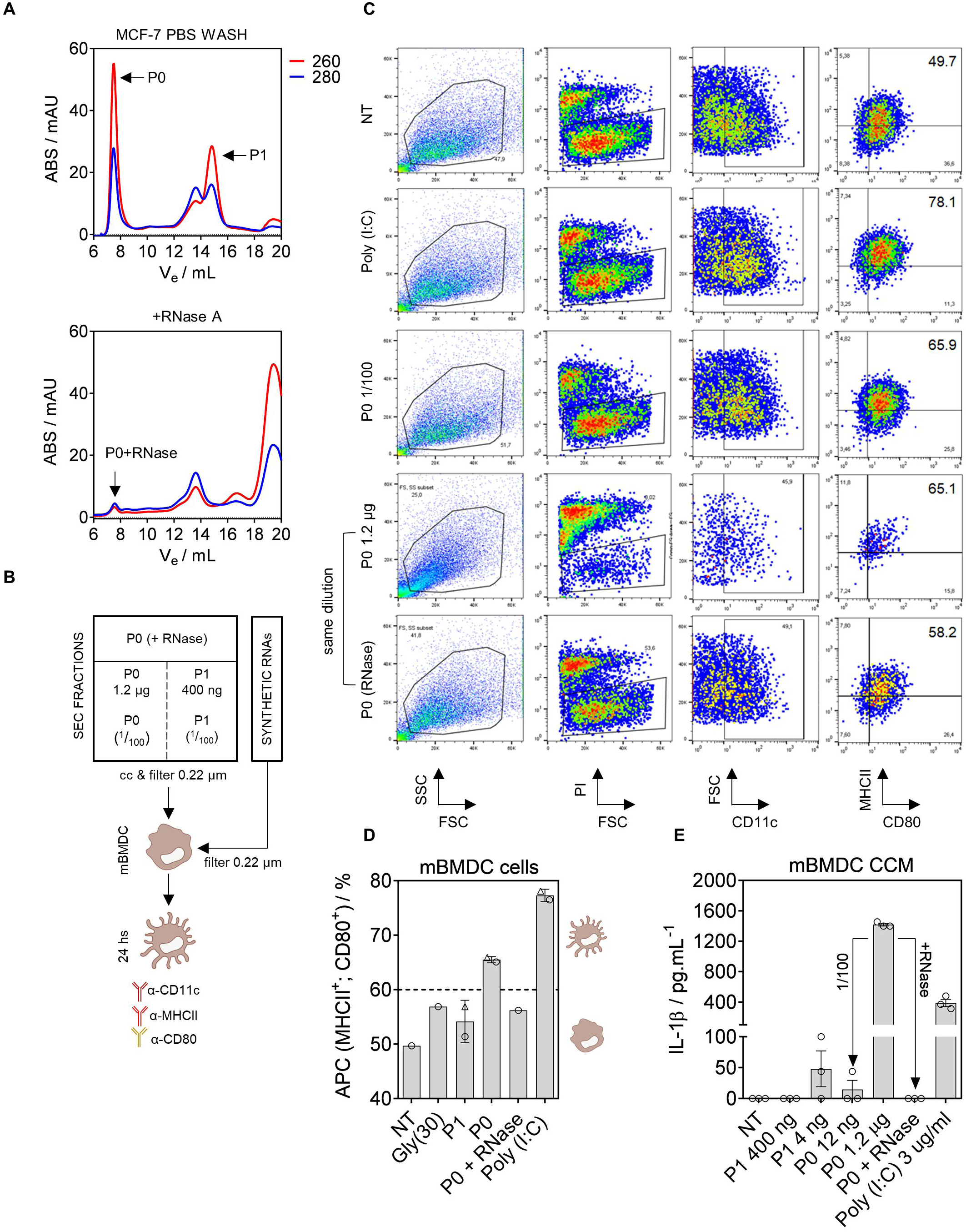
The contents of the P0 peak can trigger dendritic cell maturation in an RNA-dependent manner. A) SEC separation of the P0 and P1 peaks used for dendritic cell maturation assays. MCF-7 cells were grown in serum-free MEGM for 48 hs. The first PBS wash was discarded. Cell-conditioned PBS (t = 5 min) was concentrated and separated into two aliquots, one of which was treated with RNase A. Both samples were separated by SEC to obtain the P0 and P1 peaks (or the P0 peak from the RNase A-treated sample). B) SEC peaks were filter-sterilized and 100 µL were added to 900 µL of complete medium containing 1×10^6^ nonprimed murine bone marrow-derived dendritic cells (BMDC). The TLR3 agonist Poly (I:C) was used as a positive control. C) Flow cytometry analysis of BMDC at t = 24 hours post exposure to the P0 and P1 peaks (or synthetic RNAs). PI: propidium iodide. FSC: forward scatter. SSC: side scatter. Numbers to the right correspond to the percentage of viable (PI negative), CD11c-positive cells expressing high levels of class II MHC and CD80. D) Percentage of matured BMDC (considered as antigen-presenting cells, APC) at t = 24 hours post exposure. Triangles correspond to diluted fractions (P0: 1/100; P1: 1/100; Poly [I:C]: 1/10). E) Quantitation by ELISA of IL-1β levels in the media of BMDC analyzed by flow cytometry in the previous panel.

## DISCUSSION

A substantial fraction of exRNAs is not encapsulated inside EVs, yet the extracellular nonvesicular RNAome has not been studied in a comprehensive manner until this work. As expected, our results show that EVs and RNPs (or nonvesicular RNA complexes in general) constitute two conceptually different exRNA carriers in cell culture media that can be distinguished by their different buoyant densities and the degree of RNase protection that they confer. By inhibiting extracellular RNases, our results highlight that the nonvesicular RNA fraction is highly dynamic. This experimental approach enabled us to obtain exRNA profiles with an unprecedented level of detail and with temporal resolution. Furthermore, we succeeded in stabilizing extracellular full-length tRNAs and ribosomes, which have not been identified before outside EVs due to their susceptibility to extracellular ribonucleases. In contrast, some of their fragments were found to be highly stable and they collectively define the nonvesicular RNAome under standard conditions, especially in the presence of serum. These results have profound implications on the way we understand the mechanisms responsible for RNA release.

The presence of ribosomal aggregates in the extracellular non-EV fraction, presumably related to disomes and oligoribosomes, is further supported by the co-isolation of rRNAs, ribosomal proteins and polyA+ mRNAs from the same chromatographic fractions. Extracellular ribosomes were described in the 70’s in the blowfly *Calliphora vicina* ^57^ but subsequently linked to an experimental artefact ^58^ and have received little attention since then. However, we have demonstrated that extracellular ribosomes exist at least transiently in the media of cultured mammalian cells and possibly also in body fluids. In support of the latter, the group of Thomas Tuschl has recently optimized a modified small-RNA sequencing method that permits the identification of mRNA fragments in blood plasma or serum ^59^. Strikingly, the authors found that the distribution and length of reads mapping to mRNAs was reminiscent of ribosome profiling, suggesting that the sequenced fragments could be the footprints of ribosomes circulating in biofluids.

The biomarker potential ^60^ and the involvement of extracellular microRNAs in intercellular communication ^5^ have established a bias in the use of small RNA sequencing techniques compatible with microRNA detection to assess the RNA content of EVs. Because these techniques usually show a predominance of rRNA and tRNA-derived fragments which greatly surpass microRNAs ^30^, EVs can be considered as carriers of small RNAs between cells. However, there is an increasing amount of evidence showing that EVs actually contain more full-length ncRNAs than microRNAs or ncRNA fragments. For instance, Bioanalyzer’s peaks corresponding to intact 18S and 28S rRNAs have been identified in purified EVs ^26,61–64^, while full-length YRNAs and other ncRNAs have been identified by sequencing, RT-qPCR and/or Northern blot ^31,65^. The use of thermostable group II intron reverse transcriptases (TGIRT-seq) has allowed the identification of full-length tRNAs in EVs, which greatly outnumber tRNA-derived fragments ^53,66,67^. Our results are consistent with these reports, and clearly show the presence of tRNAs and 7SL RNAs in EVs purified by buoyant density flotation in linear iodixanol gradients. At the level of sensitivity achievable by DIG-based Northern blotting, tRNA-derived fragments were not detectable in EVs.

It is possible that different EVs derived from different subcellular compartments have different mechanisms for sorting RNAs into them ^67^. The density gradient separation method used herein was optimized to separate RNPs from EVs ^26^ rather than to discriminate between different EV subpopulations with slightly different buoyant densities ^67,68^. In any case, current evidence is sufficient to support that the most abundant intracellular RNAs are loaded and released in at least certain EV subsets ^69^.

Since RNases such as Angiogenin have been associated with EVs ^31^, it is possible that vesicle-associated RNAs may be more dynamic than previously thought. It has been reported that cancer cell-derived exosomes encapsulate Dicer and AGO2 together with pre-miRNAs and that at least some degree of cell-independent miRNA biogenesis occurs in the extracellular space ^70^. This finding is still controversial as others have not detected AGO2 or Dicer in the ultracentrifugation pellets of cell-conditioned medium ^26^ or in low-density fractions enriched in EVs ^67,71,72^. In summary, although some degree of intravesicular RNA processing is feasible, the nonvesicular extracellular fraction is intrinsically and highly dynamic while EVs tend to confer an RNase-protecting environment where less stable RNAs can persist (**Figure 6**).

**Figure 6:**
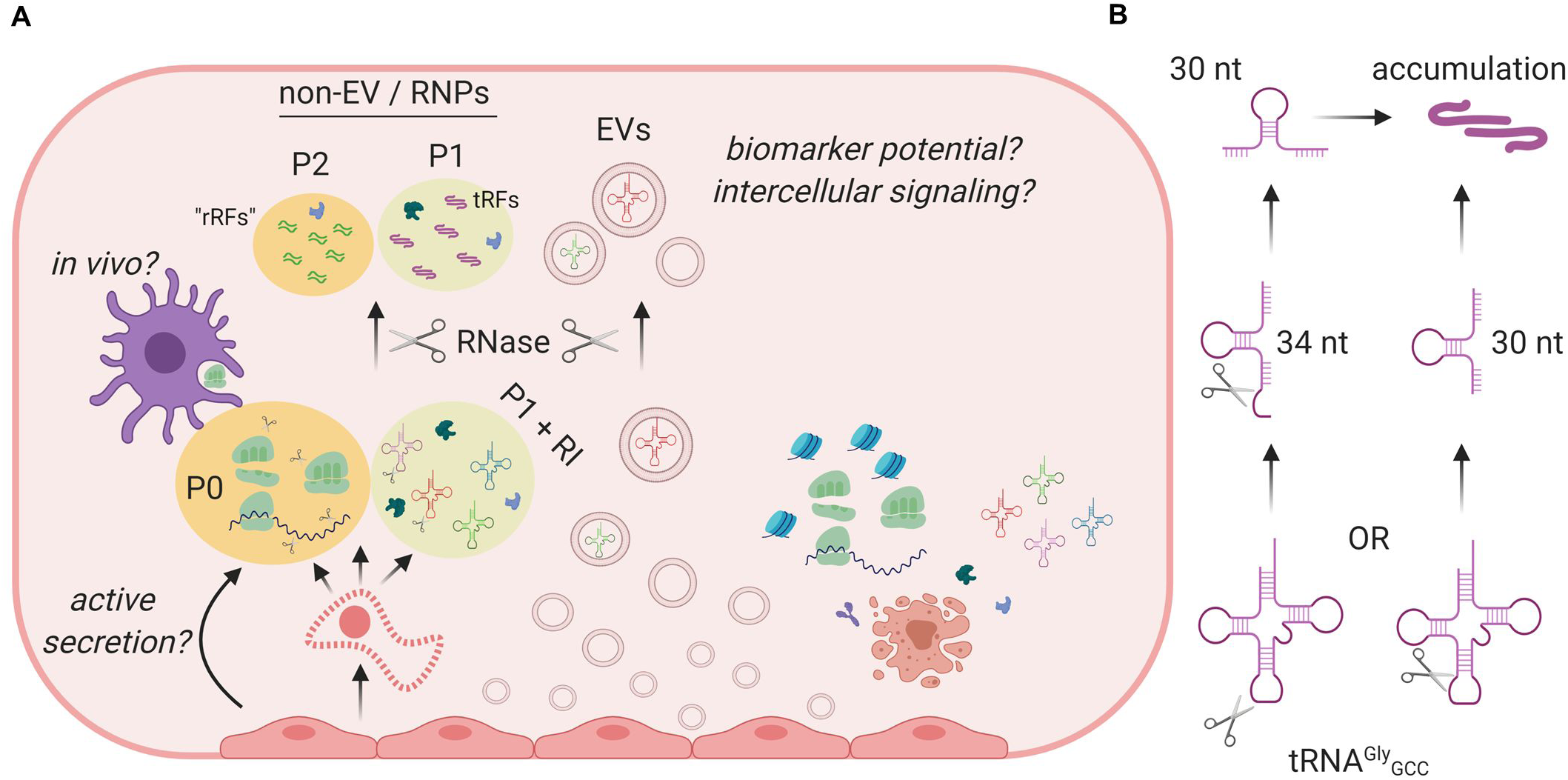
Proposed model. A) Cells in culture release tRNAs, ribosomal subunits or ribosomes to the extracellular nonvesicular space. When the CCM is analyzed by SEC, these RNAs define the P0 and P1 peaks, respectively. However, their detection is only possible after addition of RI to the medium. Regarding the mechanism responsible for the release of these RNAs, active secretion (e.g., autophagy-dependent) might contribute, but damaged or dead cells with compromised plasma membrane integrity are probably a main source of extravesicular exRNAs. Other forms of cell death can also release nucleosomes and fragmented DNA (right), although this can also occur actively by autophagy-dependent secretion. In contrast, live cells release EVs in a relatively continue fashion (center). These EVs contain ncRNAs such as tRNAs. Extracellular RNases degrade extravesicular RNAs and generate some stable fragmentation products. These products include tRNA halves, which can assemble into dimers and elute in the chromatographic P1 peak when RI is not added to the medium. We speculate that the P2 peak is composed of rRNA-derived fragments forming tightly bound dsRNAs which are not amenable to standard small RNA sequencing techniques. While full-length tRNAs and YRNAs are not detected in the non-EV fraction in the absence of RI, those which are present inside EVs are protected from degradation. Thus, EVs are probably the only source of full-length ncRNAs in RNase-rich extracellular samples. B) A diagram explaining possible biogenetic routes for extracellular, nonvesicular tRNA^Gly^_GCC_ 5’ halves.

Beyond vesicles, this work mainly focused in extravesicular exRNAs, which have received very little attention ^15^ until recently ^26^. In the past, we have compared the small RNA content between EVs and 100,000 x g supernatants of cell-conditioned medium and found that the non-EV fraction was highly enriched in 5’ tRNA halves of precisely 30 or 31 nucleotides and exclusively derived from glycine or glutamic acid tRNAs ^30^. Similar results were obtained by other groups working on primary cultures of glioblastoma ^31^. Furthermore, glycine tRNA halves are predominantly found outside EVs in serum ^34^ and are ubiquitous in many biofluids including serum, urine, saliva and bile ^32^. Altogether, these results could suggest that cells actively release specific tRNA fragments to the nonvesicular extracellular space. Herein, we have provided an alternative explanation. Enrichment of these fragments, especially when found in the non-EV fraction, can be a consequence of their differential extracellular stability rather than their preferential or selective secretion. This is further supported by the recent observation that circular RNAs, which are known to be highly stable, are increased in nearly all human biofluids when compared to matched tissues ^73^. Furthermore, we have provided evidence that the tRNA-derived fragments found outside cells are not necessarily the same as those found inside, since we have now discovered a biogenesis route for the extracellular production of tRNA halves and other related fragments. The remarkable extracellular stability of certain ncRNA fragments such as glycine 5’ tRNA halves of 30 – 31 nt (S-tRNAh) may be related to their capacity of forming homodimers ^36^. Protein binding and base modifications may confer additional layers of RNase protection.

Live cells can release a representative fraction of their cytoplasm by mechanisms such as cytoplasmic extrusion ^74^ or amphisome fusion with the plasma membrane ^26^. However, a few events of cellular damage or death might be quantitatively more important in defining exRNA profiles as has been discussed above. In support of this, it has been shown that extracellular rRNA levels correlate with extracellular lactate dehydrogenase (LDH) activity, which is widely used as a marker of cell death ^75^. Even though exRNA analysis derived from dead cells can be considered as an artifact of cell culture, there are situations where nonapoptotic, immunogenic cell death (ICD) occurs at abnormal frequencies in an organism. These situations include aging ^76^, trauma ^77^, ischemia-reperfusion injury ^78^, infectious diseases and cancer. In the latter, ICD can occur because of the hypoxic inner mass characteristic of solid tumors or following treatment with cytotoxic agents ^79^. In all cases, dying cells release intracellular components which are sensed by innate immune cells and interpreted as damage-associated molecular patterns (DAMPs). Furthermore, the therapeutic activity of several anticancer drugs eliciting ICD involves an autocrine and paracrine circuit which depends at least in part on the release of self RNAs by stressed or dying cancer cells ^80^. Because rRNAs and tRNAs are highly abundant intracellularly and they are exposed in the extracellular space in cases of damage, and considering RNAs are actively sensed by the innate immune system ^55,81^, we hypothesized that exRNA-containing nonvesicular complexes could be endowed with immunomodulatory abilities. The high turnover of these complexes as a consequence of extracellular RNases could prevent activation under physiological conditions.

As for the ribonuclease responsible for extracellular, nonvesicular tRNA cleavage, it is clearly a serum-derived ribonuclease in FBS-containing samples (probably RNase A itself). When serum is not present or highly diluted, such as after thoroughly washing cells with serum-free media or buffers, it is possible that endogenous secreted RNases could shape nonvesicular exRNA profiles. Stressed cancer cells secrete enzymes to perform some metabolic reactions in the extracellular space and then uptake the enzymatic products to fuel cellular energetics ^82^. By analogy, we are tempted to speculate that secreted RNases such as ANG could play a role in extracellular RNA metabolism, preventing the toxicity associated with its intracellular activity in nonstressed cells ^83^. Although the function of ANG in tRNA cleavage seems to be partially redundant ^51,52^, its implications in extracellular RNA cleavage under physiological conditions remains to be elucidated. Redundancy might be lower in serum-free environments as the nervous system, where several mutations in ANG have been functionally linked to neurodegenerative diseases ^84^. It is thought that mutations in ANG which impair its RNase activity will also impair angiogenesis in patients with amyotrophic lateral sclerosis (ALS) ^85^. Alternatively, ANG can confer cytoprotection against stress by the action of certain stress-induced tRNA-derived small RNAs ^86^. The possibility that ANG is involved in shaping the extracellular RNAome in cerebrospinal fluid remains unexplored. We have provided preliminary evidence suggesting an involvement of extracellular, nonvesicular RNAs or RNPs in immune surveillance. Thus, a link between mutations in ANG and deregulated extracellular RNA fragmentation patterns is feasible in diseases such as ALS whose etiology or evolution is deeply connected to inflammation ^87^.

Bacterial rRNA and tRNAs induce Toll-like receptor (TLR)-dependent DC maturation and IL-1β secretion, and are therefore considered pathogen-associated molecular patterns. However, to elicit such a response, addition of the purified RNAs with cationic lipids seems to be essential ^88^. In contrast, we have obtained high extracellular levels of IL-1β when incubating BMDC with approximately one microgram of RNA obtained from the P0 peak of MCF-7 cells (composed mainly of ribosome particles) in the absence of any transfection reagent. Strikingly, this effect was lost when incubating DCs with RNase A-pretreated P0. It remains to be elucidated whether RNA itself or any potentially associated RNA-binding proteins are responsible for these effects. Alternatively, the entire ribosome nanoparticle might be needed for efficient uptake and subsequent intracellular signaling. In any case, it is becoming increasingly clear that innate immune sensors originally thought to recognize pathogenic RNAs are used to sense damaged tissue or dying cells ^55,89,90^. It could be argued that the frailty of extracellular ribosomes might not be compatible with their capacity to elicit effective responses, especially once these particles are diluted in the extracellular space. However, DCs and distressed or damaged tumor cells are present in close contact in the tumor microenvironment ^91,92^, where soluble molecules are deeply involved in the cancer/immune cell cross-talk which ultimately determines tumor survival.

Although extracellular ribosomes are not predicted to resist extracellular RNases and are probably unrelated to recently described “exomers” ^16^, they might still achieve functionally relevant concentrations *in vivo* in extracellular microenvironments. The identification of a RNase-resistant peak (P2) derived from partial fragmentation of P0 suggests that, similarly to what we have shown for 30 – 31 nt tRNA halves, rRNA-derived fragments may accumulate in the extracellular space and their extracellular concentration may increase in situations of abnormal cell death. A new method has been recently described enabling RNA sequencing from a few microliters of human serum ^93^. With this method, almost perfect separation (average area under curve = 1 in ROC curves) between normal and breast cancer patients was possible based on rRNA or mitochondrial tRNA sequences. Because most of the serum samples were collected during or after chemotherapy, rRNA-derived fragments stably circulating in sera have the potential to alert abnormal cytotoxicity. Considering RNA-seq studies typically suppress rRNAs during library preparation, their potential as biomarkers of early onset disease in untreated patients remains an open question.

In conclusion, ribonuclease inhibition dramatically shapes extracellular RNA profiles and uncovers a population of extracellular ribosomes, tRNAs and other coding and noncoding RNAs. These RNAs, which are not protected by encapsulation inside EVs, are rapidly degraded by extracellular RNases. However, some of their fragments resist degradation and can accumulate in cell culture media and in biofluids. This dynamic view of exRNAs impacts our understanding of RNA secretion mechanisms and may offer a window to new molecules with biomarker potential. These include intrinsically stable ncRNA fragments and extracellular RNPs stabilized by addition of RI immediately upon collection of samples. The signaling potential of EV-associated and EV-free exRNAs are discussed, and the possibility that extracellular ribosomes could signal as DAMPs is presented.

## ACKNOWLEDGMENTS

JPT, MS, MH and AC are members of the Sistema Nacional de Investigadores (SNI, ANII). Partial funding for this work was received from Agencia Nacional de Investigación e Innovación, (ANII, Uruguay) [FCE_3_2018_1_148745], from Comisión Sectorial de Investigación Científica (CSIC-UdelaR, Uruguay) [MIA_PAS2019_2_99] and from the National Institutes of Health, USA [UG3CA241694, supported by the NIH Common Fund, through the Office of Strategic Coordination/Office of the NIH Director]. The authors want to thank the following core facilities at the Pasteur Institute of Montevideo: Flow cytometry (UBC), and the Analytical Biochemistry and Proteomics Unit (UByPA) for their technical assistance. TEM images were obtained in Centro Universitario Regional Este (CURE, Universidad de la República) by Álvaro Olivera with the help of Laura Fornaro. The authors also want to thank Eric Westhof (Strasbourg University), Marco Li Calzi (IPMon), Mauricio Castellano (IPMon/UdelaR), Kenneth W. Witwer (Johns Hopkins University), Paul Anderson, Shawn Lyons, Nancy Kedersha, Prakash Kharel and Vivek M. Advani (BWH/HMS) for helpful suggestions, experimental help and insightful scientific discussions.

## FIGURE LEGENDS

**Supplementary Figure 1:** data associated to Figure 1. A) Injection of synthetic RNAs of 30 nt corresponding to 5’ tiRNA^Gly^ _GCC_ (which forms RNA dimers as reported in Tosar et al. (2018) ^36^; red) and a mutant with a 25U/C substitution (which is not able to dimerize; violet) in a Superdex 200 10/300 column with PBS 1x as the mobile phase. B-C) same as Figure 1(H) and Figure 1(I) but in the P1 peak of MCF-7 CCM either treated (top) or not treated (bottom) with RI. D) Representation of SNORD49A (U49A; black)/28S rRNA (red) interaction, as depicted in snoRNABase (www-snorna.biotoul.fr). Below is the sequence with the highest number of reads. Its relative abundance is expressed as reads per million mapped reads (RPM). Its ranking in the “P1 + RI” dataset is also shown. E) Alignment of reads mapping to tRNA^Glu^ (anticodons CUC and UUC) and the genomic sequence for tRNA^Glu^_UUC_ with manual addition of the 3’ CCA sequence. “A.C”: anticodon. F) Coverage plots of sequences mapping to 28S rRNA in P0 (red), “P1 + RI” (green), “P1 – RI” (violet) and P2 (blue), either in linear scale (top) or Log2 scale (bottom).

**Supplementary Figure 2:** data associated to Figure 2. A) Same as Figure 2(B) but washing cells with MEGM instead of PBS. B) Deconvolution of chromatograms obtained by SEC analysis of PBS washes of different adherent malignant and nonmalignant cell lines, derived from different mammalian species. C) U2-OS cells before (left) and after (right) four consecutive washes with HBSS for 30 seconds.

**Supplementary Figure 3:** data associated to Figures 2 and 3. A) SEC analysis of cell-conditioned PBS of the human THP-1 cell line. Sample preparation based on protocol #5. B) Flow cytometry analysis of EG.7-OVA cells stained with propidium iodide and Annexin V-FITC. Left: nontreated cells. Center: Washed with DMEM and PBS, following protocol #5. Right: positive control for apoptotic induction with 500 µM sodium deoxycholate. C) Quantitation of the percentage of PI-negative, AnV-positive cells (left), PI-positive, AnV-positive cells (center) and PI-positive, AnV-negative cells (right) in three replicates of the experiment. *: p < 0.05. Student t test. D) Negative-stain TEM image of a ribosome-like particle in the concentrated chromatographic P0 peak from Hep G2 cells washed with PBS + RI. Scale bar: 30 nm. S: presumed small ribosomal subunit. L: presumed large ribosomal subunit. Particle dimensions: 29.0 nm x 26.2 nm. E) Density of the twelve fractions collected after high resolution 12 – 36% iodixanol gradients (related to Figure 3, G), and comparison with the reported densities in Jeppesen et al. (2019) ^26^. F) Effect of divalent cations on the P0 peak. MCF-7 cells were washed with PBS (wash #1), PBS (wash #2), PBS + 1 mM CaCl_2_ (#3) and PBS (wash #4) and all these washes were analyzed by SEC following standard procedures. G) Quantitation of the P0/P1 ratio (left), or individual P0 and P1 concentrations (obtained by deconvolution analysis of SEC chromatograms) in MCF-7 cells washed with different buffers or media. From left to right: PBS, PBS + 1mM CaCl_2_, PBS + 1mM CaCl_2_ + 1mM MgCl_2_, PBS + 1mM CaCl_2_ + 1mM MgCl_2_ + 1x nonessential amino acids (NEAA), PBS + 1x NEAA, DMEM and MEGM. *: p < 0.05; **: p < 0.01; Student t test, unpaired values. H) Quantitation of the P0 and P1 peaks when washing cells with PBS with increasing CaCl_2_ concentrations.

**Supplementary Figure 4:** data associated to Figure 4. A) Experimental conditions were similar to those used in Figure 4 A-D, but the cell-free processing step was omitted. Control lanes include RNA lysates from U2-OS cells incubated in DMEM + 10% FBS (S+) or in the same cells used for exRNA analysis (ITS, 1 hour). L: 5’ tRNA halves of 33 – 34 nt; S: 5’ tRNA halves of 30 – 31 nt. B) Comparison of SYBR gold-stained denaturing 10% PAGE gels from Figure 4, C and Supplementary Fig. 4, A. A parameter named the RNA Degradation Number (RDN) was defined as the ratio between SYBR gold intensities above and below the tRNA band. The lower the RDN (the higher extent of extracellular fragmentation), the higher the fragment-to-full-length-tRNA ratio (estimated by densitometric analysis of Northern blot bands). C) Controlled digestion of synthetic RNAs (10 nmol; 5’ phosphorylated) corresponding to 34 nt and 30 nt 5’ tRNA^Gly^_GCC_ fragments with RNase A from bovine origin (0, 0.64 and 64 pg). Reaction conditions: 5 µL, PBS 1x, 1 hour at room temperature. D) Effect of RI addition (120 U in 10 mL) in exRNA profiles from U2-OS CCM (1 hour in ITS medium or 1 – 24 hours in S+ medium). E) Isopycnic centrifugation in iodixanol gradients (12 – 36 %) of U2-OS CCM (t = 24 hs, S+) in either wild-type or ΔYBX-1 (ΔYB1) cells. Only fractions corresponding to extracellular vesicles are shown. Control lane: 1 µg intracellular RNA from U2-OS cells. F) Analysis of YRNAs (left) or some selected 5’ fragments (right) by RT-qPCR (left) or SL-RT-qPCR (right) in 100,000 x g pellets (EVs) or concentrated 100,000 x g supernatants (RNPs) of U2-OS ΔANG conditioned medium (t = 48 hs; MEGM).

**Supplementary Figure 5:** data associated to Figure 5C. Complete flow cytometry analysis in all the samples included in Figure 5, D-E. Gly (31): a synthetic single-stranded RNA of 31 nucleotide with the sequence of glycine 5’ tRNA halves.

## Notes

#### Summary of Updates

Abstract updated. Flow cyt data in Fig 5 and Fig S5 analyzed with a different software producing better visualization.

## REFERENCES

1. Heitzer, E., Haque, I. S., Roberts, C. E. S. & Speicher, M. R. Current and future perspectives of liquid biopsies in genomics-driven oncology. Nature Reviews Genetics 20, 71–88 (2019).

2. Sacher, A. G. et al. Prospective validation of rapid plasma genotyping for the detection of EGFR and kras mutations in advanced lung cancer. JAMA Oncol. 2, 1014–1022 (2016).

3. Warren, J. D. et al. Septin 9 methylated DNA is a sensitive and specific blood test for colorectal cancer. BMC Med. 9, (2011).

4. Antoury, L. et al. Analysis of extracellular mRNA in human urine reveals splice variant biomarkers of muscular dystrophies. Nat. Commun. 9, (2018).

5. Thomou, T. et al. Adipose-derived circulating miRNAs regulate gene expression in other tissues. Nature 542, 450–455 (2017).

6. Zomer, A. et al. In Vivo imaging reveals extracellular vesicle-mediated phenocopying of metastatic behavior. Cell 161, 1046–1057 (2015).

7. Valadi, H. et al. Exosome-mediated transfer of mRNAs and microRNAs is a novel mechanism of genetic exchange between cells. Nat. Cell Biol. 9, 654–659 (2007).

8. Tsui, N. B. Y., Ng, E. K. O. & Lo, Y. M. D. Stability of endogenous and added RNA in blood specimens, serum, and plasma. Clin. Chem. 48, 1647–1653 (2002).

9. Ratajczak, J. et al. Embryonic stem cell-derived microvesicles reprogram hematopoietic progenitors: evidence for horizontal transfer of mRNA and protein delivery. Leukemia 20, 847–856 (2006).

10. Skog, J. et al. Glioblastoma microvesicles transport RNA and proteins that promote tumour growth and provide diagnostic biomarkers. Nat. Cell Biol. 10, 1470–1476 (2008).

11. Tabet, F. et al. HDL-transferred microRNA-223 regulates ICAM-1 expression in endothelial cells. Nat. Commun. 5, (2014).

12. Vickers, K. C., Palmisano, B. T., Shoucri, B. M., Shamburek, R. D. & Remaley, A. T. MicroRNAs are transported in plasma and delivered to recipient cells by high-density lipoproteins. Nat. Cell Biol. 13, 423–435 (2011).

13. Arroyo, J. D. et al. Argonaute2 complexes carry a population of circulating microRNAs independent of vesicles in human plasma. Proc. Natl. Acad. Sci. U. S. A. 108, 5003–5008 (2011).

14. Turchinovich, A., Weiz, L., Langheinz, A. & Burwinkel, B. Characterization of extracellular circulating microRNA. Nucleic Acids Res. 39, 7223–7233 (2011).

15. Li, K. et al. Advances, challenges, and opportunities in extracellular RNA biology: insights from the NIH exRNA Strategic Workshop. JCI insight 3, (2018).

16. Zhang, H. et al. Identification of distinct nanoparticles and subsets of extracellular vesicles by asymmetric flow field-flow fractionation. Nat. Cell Biol. 20, 332–343 (2018).

17. Zhang, Q. et al. Transfer of Functional Cargo in Exomeres. Cell Rep. 27, 940-954.e6 (2019).

18. Tkach, M. & Théry, C. Communication by Extracellular Vesicles: Where We Are and Where We Need to Go. Cell 164, 1226–1232 (2016).

19. Cai, Q. et al. Plants send small RNAs in extracellular vesicles to fungal pathogen to silence virulence genes. Science 360, 1126–1129 (2018).

20. Ostrowski, M. et al. Rab27a and Rab27b control different steps of the exosome secretion pathway. Nat. Cell Biol. 12, 19–30 (2010).

21. Trajkovic, K. et al. Ceramide triggers budding of exosome vesicles into multivesicular endosomes. Science (80-.). 319, 1244–1247 (2008).

22. Hoshino, A. et al. Tumour exosome integrins determine organotropic metastasis. Nature 527, 329–335 (2015).

23. Kamerkar, S. et al. Exosomes facilitate therapeutic targeting of oncogenic KRAS in pancreatic cancer. Nature 546, 498–503 (2017).

24. Elahi, F. M., Farwell, D. G., Nolta, J. A. & Anderson, J. D. Preclinical Translation of Exosomes Derived from Mesenchymal Stem/Stromal Cells. Stem Cells (2019). doi: 10.1002/stem.3061

25. Turchinovich, A., Tonevitsky, A. G. & Burwinkel, B. Extracellular miRNA: A Collision of Two Paradigms. Trends in Biochemical Sciences 41, 883–892 (2016).

26. Jeppesen, D. K. et al. Reassessment of Exosome Composition. Cell 177, 428-445.e18 (2019).

27. Snyder, M. W., Kircher, M., Hill, A. J., Daza, R. M. & Shendure, J. Cell-free DNA Comprises an in Vivo Nucleosome Footprint that Informs Its Tissues-Of-Origin. Cell 164, 57–68 (2016).

28. Ulz, P. et al. Inferring expressed genes by whole-genome sequencing of plasma DNA. Nat. Genet. 48, 1273–1278 (2016).

29. Murillo, O. D. et al. exRNA Atlas Analysis Reveals Distinct Extracellular RNA Cargo Types and Their Carriers Present across Human Biofluids. Cell 177, 463-477.e15 (2019).

30. Tosar, J. P. et al. Assessment of small RNA sorting into different extracellular fractions revealed by high-throughput sequencing of breast cell lines. Nucleic Acids Res. 43, 5601–5616 (2015).

31. Wei, Z. et al. Coding and noncoding landscape of extracellular RNA released by human glioma stem cells. Nat. Commun. 8, 1145 (2017).

32. Srinivasan, S. et al. Small RNA Sequencing across Diverse Biofluids Identifies Optimal Methods for exRNA Isolation. Cell 177, 446-462.e16 (2019).

33. Godoy, P. M. et al. Large Differences in Small RNA Composition Between Human Biofluids. Cell Rep. 25, 1346–1358 (2018).

34. Dhahbi, J. M. et al. 5’ tRNA halves are present as abundant complexes in serum, concentrated in blood cells, and modulated by aging and calorie restriction. BMC Genomics 14, 298 (2013).

35. Zhang, Y. et al. Identification and characterization of an ancient class of small RNAs enriched in serum associating with active infection. Journal of Molecular Cell Biology 6, 172–174 (2014).

36. Tosar, J. P. et al. Dimerization confers increased stability to nucleases in 5’ halves from glycine and glutamic acid tRNAs. Nucleic Acids Res. 46, 9081–9093 (2018).

37. Fu, H. et al. Stress induces tRNA cleavage by angiogenin in mammalian cells. FEBS Lett. 583, 437–442 (2009).

38. Yamasaki, S., Ivanov, P., Hu, G.-F. & Anderson, P. Angiogenin cleaves tRNA and promotes stress-induced translational repression. J. Cell Biol. 185, 35–42 (2009).

39. Thompson, D. M., Lu, C., Green, P. J. & Parker, R. tRNA cleavage is a conserved response to oxidative stress in eukaryotes. RNA 14, 2095–2103 (2008).

40. Segovia, M., Cuturi, M. C. & Hill, M. Preparation of mouse bone marrow-derived dendritic cells with immunoregulatory properties. Methods Mol. Biol. 677, 161–168 (2011).

41. Cozen, A. E. et al. ARM-seq: AlkB-facilitated RNA methylation sequencing reveals a complex landscape of modified tRNA fragments. Nat. Methods 12, 879–884 (2015).

42. Gogakos, T. et al. Characterizing Expression and Processing of Precursor and Mature Human tRNAs by Hydro-tRNAseq and PAR-CLIP. Cell Rep. 20, 1463–1475 (2017).

43. Torres, A. G., Reina, O., Attolini, C. S. O. & De Pouplana, L. R. Differential expression of human tRNA genes drives the abundance of tRNA-derived fragments. Proc. Natl. Acad. Sci. U. S. A. 116, 8451–8456 (2019).

44. Zheng, G. et al. Efficient and quantitative high-throughput tRNA sequencing. Nat. Methods 12, 835–837 (2015).

45. Pakos□Zebrucka, K. et al. The integrated stress response. EMBO Rep. 17, 1374–1395 (2016).

46. Ouyang, D. Y. et al. Autophagy is differentially induced in prostate cancer LNCaP, DU145 and PC-3 cells via distinct splicing profiles of ATG5. Autophagy 9, 20–32 (2013).

47. Wettenhall, R. E. & Wool, I. G. Reassociation of eukaryotic ribosomal subunits. Kinetics of reassociation; role of deacylated transfer ribonucleic acid; effect of cycloheximide. J. Biol. Chem. 247, 7201–6 (1972).

48. Nolan, R. D. & Arnstein, H. R. V. The Dissociation of Rabbit Reticulocyte Ribosomes with EDTA and the Location of Messenger Ribonucleic Acid. Eur. J. Biochem. 9, 445–450 (1969).

49. Yoshikawa, H. et al. Efficient analysis of mammalian polysomes in cells and tissues using ribo mega-SEC. Elife 7, (2018).

50. Gardiner, C. et al. Techniques used for the isolation and characterization of extracellular vesicles: Results of a worldwide survey. J. Extracell. Vesicles 5, (2016).

51. Akiyama, Y. et al. Multiple ribonuclease A family members cleave transfer RNAs in response to stress. bioRxiv 811174 (2019). doi: 10.1101/811174

52. Su, Z., Kuscu, C., Malik, A., Shibata, E. & Dutta, A. Angiogenin generates specific stress-induced tRNA halves and is not involved in tRF-3-mediated gene silencing. J. Biol. Chem. jbc.RA119.009272 (2019). doi: 10.1074/jbc.RA119.009272

53. Shurtleff, M. J. et al. Broad role for YBX1 in defining the small noncoding RNA composition of exosomes. Proc. Natl. Acad. Sci. U. S. A. 114, E8987–E8995 (2017).

54. Buck, A. H. et al. Exosomes secreted by nematode parasites transfer small RNAs to mammalian cells and modulate innate immunity. Nat. Commun. 5, 5488 (2014).

55. Gong, T., Liu, L., Jiang, W. & Zhou, R. DAMP-sensing receptors in sterile inflammation and inflammatory diseases. Nat. Rev. Immunol. (2019). doi: 10.1038/s41577-019-0215-7

56. Banchereau, J. & Steinman, R. M. Dendritic cells and the control of immunity. Nature 392, 245–252 (1998).

57. Sridhara, S. & Levenbook, L. Extracellular ribosomes during metamorphosis in the blowfly Calliphora erythrocephala. Biochem. Biophys. Res. Commun. 53, 1253–9 (1973).

58. Levenbook, L., Sridhara, S. & Lambertsson, A. Extracellular ribosomes during metamorphosis of the blowfly Calliphora vicina R. D. -- A reappraisal of their authenticity. Biochem. Biophys. Res. Commun. 70, 15–21 (1976).

59. Akat, K. M. et al. Detection of circulating extracellular mRNAs by modified small-RNA-sequencing analysis. JCI Insight 4, 127317 (2019).

60. Yokoi, A. et al. Integrated extracellular microRNA profiling for ovarian cancer screening. Nat. Commun. 9, (2018).

61. Lunavat, T. R. et al. BRAFV600 inhibition alters the microRNA cargo in the vesicular secretome of malignant melanoma cells. Proc. Natl. Acad. Sci. U. S. A. 114, E5930–E5939 (2017).

62. Lunavat, T. R. et al. Small RNA deep sequencing discriminates subsets of extracellular vesicles released by melanoma cells – Evidence of unique microRNA cargos. RNA Biol. 12, 810–823 (2015).

63. Lässer, C. et al. Two distinct extracellular RNA signatures released by a single cell type identified by microarray and next-generation sequencing. RNA Biol. 14, 58–72 (2017).

64. Chiou, N. T., Kageyama, R. & Ansel, K. M. Selective Export into Extracellular Vesicles and Function of tRNA Fragments during T Cell Activation. Cell Rep. 25, 3356-3370.e4 (2018).

65. Driedonks, T. A. P. et al. Immune stimuli shape the small non-coding transcriptome of extracellular vesicles released by dendritic cells. Cell. Mol. Life Sci. 75, 3857–3875 (2018).

66. Qin, Y. et al. High-throughput sequencing of human plasma RNA by using thermostable group II intron reverse transcriptases. RNA 22, 111–128 (2016).

67. Temoche-Diaz, M. M. et al. Distinct mechanisms of microRNA sorting into cancer cell-derived extracellular vesicle subtypes. Elife 8, e47544 (2019).

68. Kowal, J. et al. Proteomic comparison defines novel markers to characterize heterogeneous populations of extracellular vesicle subtypes. Proc. Natl. Acad. Sci. U. S. A. 113, E968–E977 (2016).

69. Gámbaro, F. et al. Stable tRNA halves can be sorted into extracellular vesicles and delivered to recipient cells in a concentration-dependent manner. RNA Biol. in press, 1–15 (2019).

70. Melo, S. A. et al. Cancer exosomes perform cell-independent microRNA biogenesis and promote tumorigenesis. Cancer Cell 26, 707–721 (2014).

71. Shurtleff, M. J., Temoche-Diaz, M. M., Karfilis, K. V, Ri, S. & Schekman, R. Y-box protein 1 is required to sort microRNAs into exosomes in cells and in a cell-free reaction. Elife 5, e19276 (2016).

72. Van Deun, J. et al. The impact of disparate isolation methods for extracellular vesicles on downstream RNA profiling. J. Extracell. Vesicles 3, 24858 (2014).

73. Hulstaert, E. et al. Charting extracellular transcriptomes in The Human Biofluid RNA Atlas. bioRxiv 823369 (2019). doi: 10.1101/823369

74. Lee, K.-Z. et al. Enterocyte Purge and Rapid Recovery Is a Resilience Reaction of the Gut Epithelium to Pore-Forming Toxin Attack. Cell Host Microbe 20, 716–730 (2016).

75. Böttcher, K., Wenzel, A. & Warnecke, J. M. Investigation of the origin of extracellular RNA in human cell culture. Ann. N. Y. Acad. Sci. 1075, 50–6 (2006).

76. Franceschi, C., Garagnani, P., Vitale, G., Capri, M. & Salvioli, S. Inflammaging and ‘Garb-aging’. Trends Endocrinol. Metab. 28, 199–212 (2017).

77. Zhang, Q. et al. Circulating mitochondrial DAMPs cause inflammatory responses to injury. Nature 464, 104–107 (2010).

78. Slegtenhorst, B. R., Dor, F. J. M. F., Rodriguez, H., Voskuil, F. J. & Tullius, S. G. Ischemia/Reperfusion Injury and its Consequences on Immunity and Inflammation. Curr. Transplant. Reports 1, 147–154 (2014).

79. Galluzzi, L., Buqué, A., Kepp, O., Zitvogel, L. & Kroemer, G. Immunogenic cell death in cancer and infectious disease. Nature Reviews Immunology 17, 97–111 (2017).

80. Sistigu, A. et al. Cancer cell–autonomous contribution of type I interferon signaling to the efficacy of chemotherapy. Nat. Med. 20, 1301–1309 (2014).

81. Dhir, A. et al. Mitochondrial double-stranded RNA triggers antiviral signalling in humans. Nature 560, 238–242 (2018).

82. Loo, J. M. et al. Extracellular metabolic energetics can promote cancer progression. Cell 160, 393–406 (2015).

83. Thomas, S. P., Hoang, T. T., Ressler, V. T. & Raines, R. T. Human angiogenin is a potent cytotoxin in the absence of ribonuclease inhibitor. RNA 24, 1018–1027 (2018).

84. van Es, M. A. et al. Angiogenin variants in Parkinson disease and amyotrophic lateral sclerosis. Ann. Neurol. 70, 964–73 (2011).

85. Lambrechts, D., Lafuste, P., Carmeliet, P. & Conway, E. M. Another angiogenic gene linked to amyotrophic lateral sclerosis. Trends in Molecular Medicine 12, 345–347 (2006).

86. Ivanov, P. et al. G-quadruplex structures contribute to the neuroprotective effects of angiogenin-induced tRNA fragments. Proc. Natl. Acad. Sci. U. S. A. 111, 18201–18206 (2014).

87. Trias, E. et al. Mast cells and neutrophils mediate peripheral motor pathway degeneration in ALS. JCI insight 3, (2018).

88. Sha, W. et al. Human NLRP3 Inflammasome senses multiple types of bacterial RNAs. Proc. Natl. Acad. Sci. U. S. A. 111, 16059–16064 (2014).

89. Cavassani, K. A. et al. TLR3 is an endogenous sensor of tissue necrosis during acute inflammatory events. J. Exp. Med. 205, 2609–2621 (2008).

90. Feng, Y. et al. Cardiac RNA induces inflammatory responses in cardiomyocytes and immune cells via Toll-like receptor 7 signaling. J. Biol. Chem. 290, 26688–26698 (2015).

91. Veglia, F. & Gabrilovich, D. I. Dendritic cells in cancer: the role revisited. Current Opinion in Immunology 45, 43–51 (2017).

92. Tran Janco, J. M., Lamichhane, P., Karyampudi, L. & Knutson, K. L. Tumor-Infiltrating Dendritic Cells in Cancer Pathogenesis. J. Immunol. 194, 2985–2991 (2015).

93. Zhou, Z. et al. Extracellular RNA in a single droplet of human serum reflects physiologic and disease states. Proc. Natl. Acad. Sci. 201908252 (2019). doi: 10.1073/pnas.1908252116

