## Supplementary material for "Fragmentation of extracellular ribosomes and tRNAs shapes the extracellular RNAome": SI Materials and Methods

[1]Analytical Biochemistry Unit. Nuclear Research Center. Faculty of Science. Universidad de la República, Uruguay. [2]Functional Genomics Unit, Institut Pasteur de Montevideo, Uruguay. [3]Division of Rheumatology, Immunology and Allergy, Brigham and Women's Hospital, Boston, MA, USA. Department of Medicine, Harvard Medical School, Boston, MA, USA. [4]Laboratory of Immunoregulation and Inflammation, Institut Pasteur de Montevideo, Uruguay. Immunobiology Department, Faculty of Medicine, Universidad de la República, Uruguay. [5]Molecular Virology Laboratory, Nuclear Research Center. Faculty of Science. Universidad de la República, Uruguay. [6]The Broad Institute of Harvard and M.I.T., Cambridge, MA, USA. [7]Department of Medicine, University Hospital, Universidad de la República, Uruguay.

### SUPPLEMENTARY METHODS

#### Reagents

RI (ribonuclease inhibitor, murine; 40U / µL) was purchased from New England Biolabs. Sterile Phosphate buffered saline, PBS (10x DPBS for chromatographic separations, 1x DPBS with or without calcium and magnesium for cell culture), DMEM, trypsin–EDTA solution, fetal bovine serum (FBS), 100x Insulin-Transferrin-Selenium solution (ITS) and nuclease-free distilled water were obtained from Gibco. Hank’s Balanced Salt Solution (HBSS, no phenol red), either with or without calcium and magnesium were obtained from Corning as 1x sterile solutions. Trizol and Trizol LS reagents were from Invitrogen (Thermo). The 60% Optiprep solution used to prepare iodixanol gradients was obtained from Sigma.

#### Cell lines

Cell lines were obtained from ATCC. Creation and characterization of U2-OS ΔANG and U2-OS ΔRNH1 cells was described in Akiyama et al. 2019 (doi: 10.1101/811174). The gene-edited cells were clonally selected and genotyped.

#### Antibodies

The following primary antibodies were used for Western blot: anti-Puromycin (Millipore, cat # MABE343, clone 12D10); anti-RPS6 (Santa Cruz Biotechnology; cat # sc-74459; clone C-8); anti-RPS23 (Santa Cruz Biotechnology; cat # sc-100837; clone SJ-K2); anti-RPL7a (Cell Signaling; cat # 2415; clone E109); anti-CD63 (BD Biosciences; cat # 556019; clone H5C6); anti-CD9 (Millipore; cat # CBL162; clone MM2/57); anti-CD81 (R&D; cat # MAB4615; clone 454720); anti-eIF2α (Cell Signaling: cat # 9722); anti-(p)eIF2α (Ser51; Cell Signaling; cat # 9721). Antibodies for flow cytometry where from BD and directed to mouse proteins CD11c (clone HL3), CD80 (clone 16-10A1), I-Ab (clone AF6-120.1).

#### Primers and RT-qPCR

In figure 1J: cDNA was obtained with SuperScript II (Thermo) using random hexamers and following manufacturer’s instructions. For glycine 5’ halves a gene-specific primer (GSP) was used (the sample was heated at 65$^{o}$C, following by primer annealing and extension at 42$^{o}$C). Input material: 2 µL of each chromatographic fraction. Quantitative real-time PCR was performed with a Kapa SYBR Fast qPCR Master Mix (2x) from Kapa Biosystems.

tRNA$^{Gly}$ 5' halves (RT GSP): TGCCATCCACCACCCTGTTGCTGTAGCGAGAATT
tRNA$^{Gly}$ 5' halves (F-primer): ccCCGCATTGGTGGTTCAGTGGTA
tRNA$^{Gly}$ 5' halves (R-primer): TCCACCACCCTGTTGCTGTA

28S rRNA (position 310; F-primer): GGGTGGTAAACTCCATCTAAGG
28S rRNA (position 310; R-primer): GCCCTCTTGAACTCTCTCTTC
28S rRNA (position 3744; F-primer): GTAAACGGCGGGAGTAACTATG
28S rRNA (position 3744; R-primer): GACAGTGGGAATCTCGTTCATC
18S rRNA (position 442; F-primer): CTGAGAAACGGCTACCACATC
18S rRNA (position 442; R-primer): GCCTCGAAAGAGTCCTGTATTG
5.8S rRNA (position 25; F-primer): CTCGTGCGTCGATGAAGAA
5.8S rRNA (position 25; R-primer): TCGAAGTGTCGATGATCAATGT
5.8S rRNA (position 2; F-primer): ACTCTTAGCGGTGGATCACT
5.8S rRNA (position 2; R-primer): GATGATCAATGTGTCCTGCAATTC

In Figure 3E: Chromatographic fractions corresponding to the P0 peak from BJ cells were concentrated to 50 µL by ultrafiltration (Vivaspin 500; MWCO 5 kDa). One microliter was used as input. cDNA was obtained with SuperScript II (Thermo) using an oligo$(dT)_{18}$ primer and following manufacturer's instructions. End-point PCR was performed with recombinant Taq DNA Polymerase (Thermo) for 35 cycles. Annealing temperature: 58$^{o}$C. Expected amplicon length: 731 bp ( > 2200 bp from predicted mRNA 3' end)

HSP90B1 (F-primer): GGTGTAGGAATGACCAGAGAAG
HSP90B1 (R-primer): GGAGCAGATGTGGGTACAAATA

In Supplementary Figure S4, F: cDNA for full-length YRNA analysis was obtained with SuperScript II (Thermo) using random hexamers and following manufacturer's instructions. cDNA for YRNA fragments was obtained based on the stem-loop RT-qPCR method which was described in detail in our previous publication (Tosar et al. 2018; *Nucleic Acids Research* 46, 9081-9093).

Full-length YRNAs (conventional RT with random hexamers):

RNA Y1 (F-primer): TGGTCCGAAGGTAGTGAGTTA
RNA Y1 (R-primer): GTCAAGTGCAGTAGTGAGAAGG
RNA Y3 (F-primer): CCGAGTGCAGTGGTGTTTA
RNA Y3 (R-primer): AGGGCTAGTCAAGTGAAGCAG

RNA Y4 (F-primer): GTCCGATGGTAGTGGGTTATC
RNA Y4 (R-primer): AAAGCCAGTCAAATTTAGCAGT
RNA Y5 (F-primer): GTCCGAGTGTTGTGGGTTATT
RNA Y5 (R-primer): ACAGCAAGCTAGTCAAGCG

YRNA fragments (SL-RT-qPCR; as described in Tosar et al. 2018):

Stem-loop RT primer ("X" denotes assay-specific 3' overhangs):
GTCGTATCCAGTGCAGGGTCCGAGGTATTCGCACTGGATACGACXXXXXX
RNA Y1 (5' fragment; 31 nt; 3' overhang): ATTGAG
RNA Y4 (5' fragment; 32 nt; 3' overhang): AGTTCT
RNA Y5 (5' fragment; 32 nt; 3' overhang): CTTAAC
miR-21-5p (3' overhang): GTCAAC
RNA Y1 (5' fragment; 31 nt; F-primer): caagTGGTCCGAAGGTAGTGAGT
RNA Y4 (5' fragment; 32 nt; F-primer): tgGTCCGATGGTAGTGGGTT
RNA Y5 (5' fragment; 32 nt; F-primer): agttgGTCCGAGTGTTGTGG
miR-21-5p (F-primer): gccccgTAGCTTATCAGACTGATGT
Universal reverse primer: GTGCAGGGTCCGAGGT

All primers were obtained from Integrated DNA Technologies (IDT, USA).

**Sequencing data analysis**

After sample demultiplexing and adapter trimming (only sequences >15 bases which contained an identifiable 3′ adaptor were analyzed), FastQ files containing sequencing information were mapped to the human genome (hg19) with Bowtie, allowing one mismatch. Mapped reads were collapsed to unique sequences with their associated count numbers and aligned with Lastz (one mismatch allowance) to manually curated reference libraries containing unambiguous sequences corresponding to mature human miRNAs downloaded from miRBase (http://www.mirbase.org/) release 20; mature human tRNAs downloaded from the genomic tRNA database (http://gtrnadb.ucsc.edu/); and mature human ribosomal RNA, small nuclear RNA, small nucleolar RNA, vault RNA and YRNA sequences downloaded from GenBank (http://www.ncbi.nlm.nih.gov/nuccore/). Only one annotation per read was kept, prioritizing sense alignments with no mismatches on the entire query sequence. Since the Lastz algorithm uses seeds with a minimum length of 19, the size range of the RNAs under study started at 19 nt. Sequences annotated with more than one tag were manually resolved by considering the presence/absence of mismatches, length of the query sequence and abundance-based likelihood. The relative abundance of each unique sequence was expressed as reads per million (RPM) mapped reads by dividing its absolute count number by the total amount of mapped reads in the data set and multiplying by a million. Subsequent analysis was performed with in-house ad-hoc scripts.

Data was submitted to NCBI's small read archive (SRA) under the BioProject ID: PRJNA454316. Note that the P1 and P2 peaks are denoted as H and L, respectively.

**Proteomic analysis**

Fractions corresponding to selected chromatographic peaks (0.5 mL) were treated with 640 ng RNase A and incubated for 1 h at 37$^{o}$C. The samples were then concentrated by ultrafiltration (Vivaspin 500, MWCO 5 kDa) to 50 µL by performing successive dilutions with PBS in order to remove degraded RNAs. After this, 1 µL of sequencing grade modified Trypsin (Promega) was added and incubated overnight at 37$^{o}$C. Peptides were then purified using a C18 ZipTip (Merck Millipore) following manufacturer's instructions. Eluted peptides were dried in a SpeedVac and resuspended in 10 µL of 0.1 % formic acid. Each sample was injected into a nano-HPLC system (EASY-nLC 1000, Thermo Scientific) fitted with a reverse-phase column (EASY-Spray column, 150 mm × 50 µm, C18, 2 µm, Thermo Scientific). Peptides were separated on a linear gradient of solvent B (50% ACN : 0.1% formic acid (v/v)) from 4% to 55% in 60 min.

Peptide analysis was carried out in a LTQ Velos nano-ESI linear ion trap instrument (Thermo Scientific) set in a data-dependent acquisition mode using a dynamic exclusion list. NSI-source parameters were set as follows: spray voltage (kV): 2.2 and 300 °C capillary temperature. Mass analysis was performed with Xcalibur 2.1 in two steps: (1) acquisition of full MS scan in the positive ion mode with m/z between 400 and 1200 Da, (2) CID fragmentation of the ten most intense ions with the following parameters; normalized collision energy: 35, activation Q: 0.25; activation time: 15 ms.

Proteomic data analysis was performed with the program PatternLab for Proteomics (version 3.2.0.3) and using a database containing the *Mus musculus* proteome downloaded from Uniprot (2017/12/21) and 127 of the most common mass spectrometry contaminants. The Comet search engine was set as follows: tryptic peptides; oxidation of methionine as variable modification and 800 ppm of tolerance from the measured precursor m/z. Peptide spectrum matches were filtered using the Search Engine Processor (SEPro) using the following parameters; acceptable FDR: 1% at the protein level and a minimum of two peptides per protein.

**Northern blotting**

RNA samples were run on 10% TBE-urea polyacrylamide gels (ThermoFisher Scientific), transferred to positively charged nylon membranes (Roche). The membranes were cross-linked by UV irradiation. After cross-linking, the membranes were hybridized overnight at 40°C with digoxigenin (DIG)-labeled DNA probes in DIG Easy Hyb solution (Roche). After low stringency washes (washing twice with 2× SSC/0.1% SDS at room temperature) and a high stringency wash (1× SSC/0.1% SDS at 40ºC), the membranes were blocked in blocking reagent (Roche) for 30 min at room temperature, probed with alkaline phosphatase-labeled anti-digoxigenin antibody (Roche) for 30 min, and washed with 1x TBS-T. Signals were visualized with CDP-Star ready-to-use (Roche) and detected using ChemiDoc imaging system (BioRad) according to the manufacturer's instructions. Oligonucleotide probes were synthesized by IDT. DIG-labeled probes were prepared using the DIG Oligonucleotide tailing kit (2nd generation; Roche) according to the manufacturer's instructions. The sequences of the probes were as follows:

probe for 5'-$tRNA^{Lys}_{UUU}$ : 5' CTGATGCTCTACCGACTGAGCTATCCGGGC 3';
probe for 5'-$tRNA^{iMet}_{CAU}$ : 5' CTTCCGCTGCGCCACTCTGCT 3';
probe for 5'-$tRNA^{Gly}_{GCC}$ : 5' CTACCACTGAACCACCCATGC 3';
probe for 3'-$tRNA^{Gly}_{GCC}$ : 5' GCCGGGAATCGAACCCGGGCCTCCCGCG 3';
probe for 7SL RNA: 5' CACTACAGCCCAGAACTCCTGGACT 3'.

**Transmission electron microscopy**

Two T75 flasks containing Hep G2 cells grown in S+ at 80% confluency were washed twice with serum-free DMEM, once with PBS, and once with PBS + 40U RI. This last wash was concentrated and subjected to size exclusion chromatography as previosuly described. The fractions corresponding to the P0 peak were concentrated to 20 µL by ultrafiltration and frozen. Once thawed, they were incubated on carbon-coated grids for 1.5 min. The grids were washed twice in miliQ water for 30 seconds, and placed inverted on top of a droplet containing 1% phosphotungstic acid (twice). Grids were dried and imaged in a JEOL JEM-2100 electron microscope at 200 kV.

**Sample preparation for dendritic cell maturation assays**

Two T75 flasks containing MCF-7 cells were grown in MEGM for 48 hours (80% confluency) and then washed with PBS + 40U RI for 30 seconds. The cell-conditioned PBS was separated into two aliquotes. One aliquot was treated with 320 ng RNase A and incubated at $37^{o}$C for 30 min. Both aliquotes were then subjected to SEC in order to obtain the following fractions/samples: P0, RNase-treated P0 and P1. Each of these fractions were concentrated to 100 µL by ultrafiltration (MWCO 5 kDa) and filter-sterilized in the cell culture room. P0 and P1 were also diluted 100-fold with sterile PBS. All samples (100 µL) were added directly to the media of 1 x $10^6$ BMDCs grown in 900 µL of complete media (RPMI + 10% FBS, 0.05 mM 2-mercaptoethanol, 2mM L-glutamine, 1mM sodium pyruvate, 1% HEPES, 100 U/mL penicillin, 0.1 mg/mL streptomycin ; 6-well plates) and incubated for 24 hours at $37^{o}$C and 5% $CO_2$. Control wells contained: 100 µL PBS (NT; nontreated cells), synthetic $tRNA^{Gly}_{GCC}$ 5' halves or a mutated version of this RNA ($25^{U/C}$; both at 10 µg / mL final concentration), or the synthetic dsRNA analogue Poly(I:C) (Invivogen) at either 30 µg / mL or 3 µg / mL, diluted in PBS.

At that time, cells were harvested and resuspended in PSA (PBS, 0.2% fetal bovine serum, 0.1 sodium azide) to stain them with antibodies against CD11c, CD80 and I-Ab for 20 minutes on light-protected ice. After two washes with PBS, propidium iodide was added. The samples were acquired in the CyAN ADP Analyzer (Dako) and the data analyzed with the FlowJo vX.0.7 software (FlowJo, LLC)
