## Supplementary figures and images for "Fragmentation of extracellular ribosomes and tRNAs shapes the extracellular RNAome"

### Supplementary Fig. 1

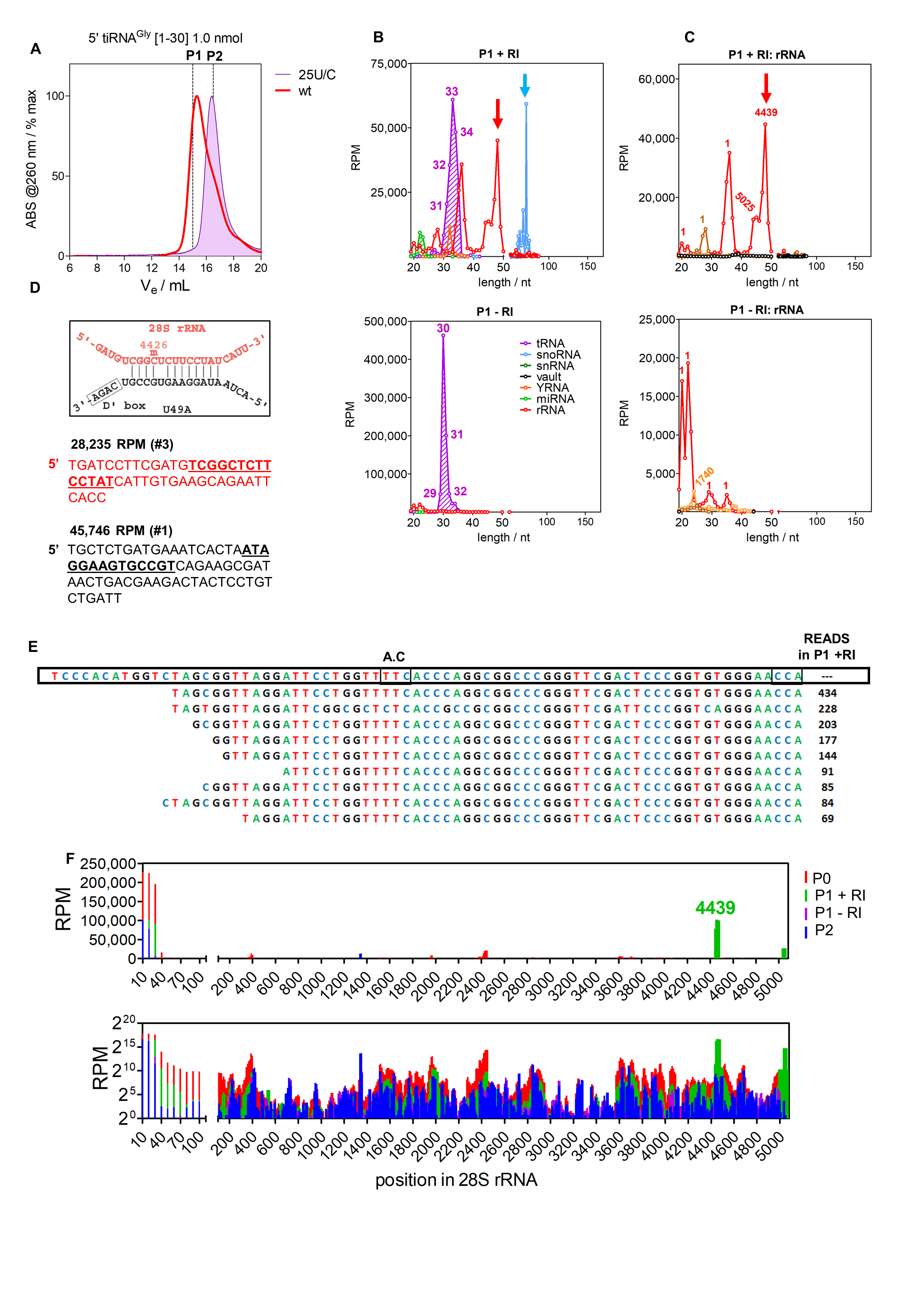

### Supplementary Fig. 2

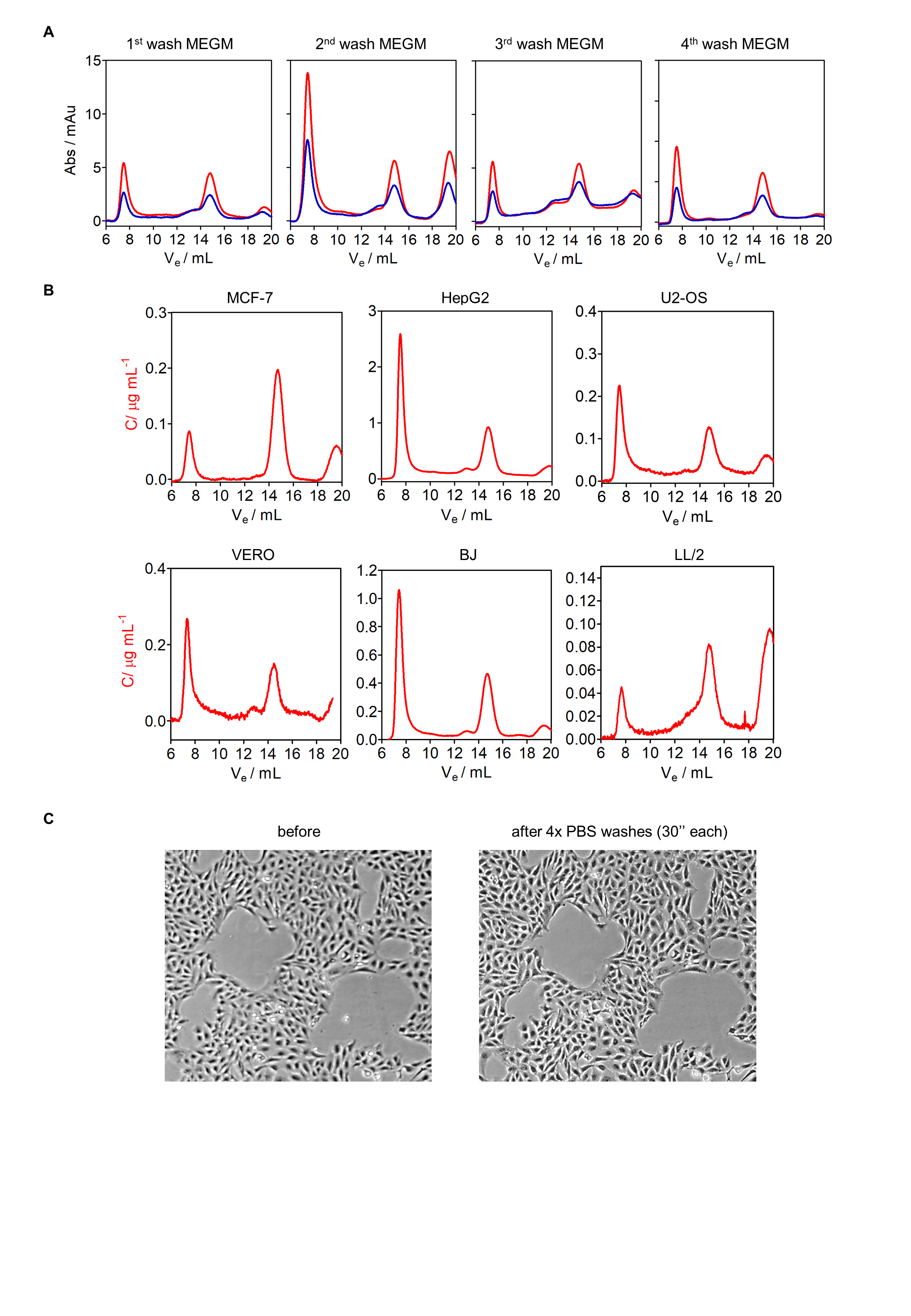

### Supplementary Fig. 3

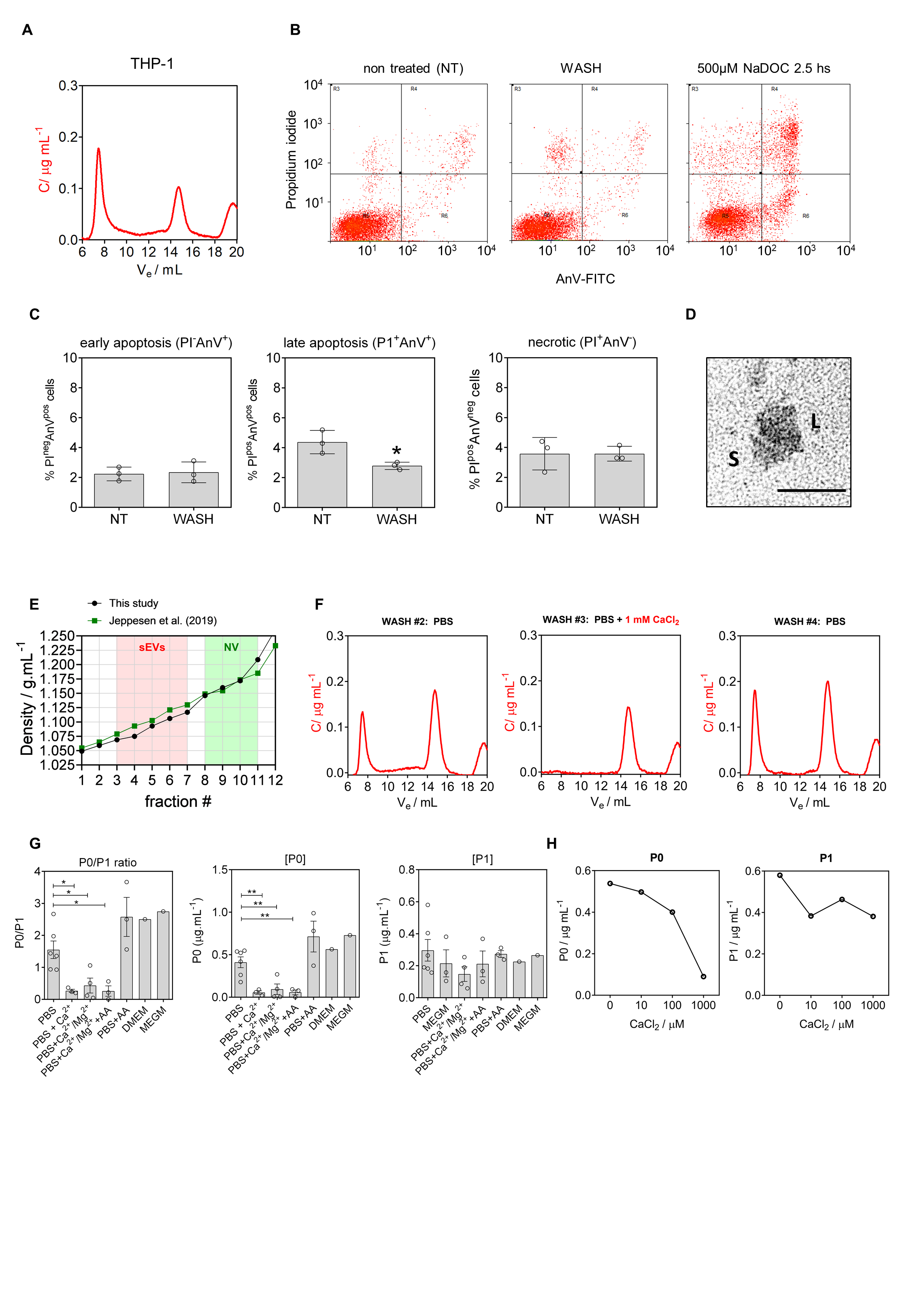

### Supplementary Fig. 4

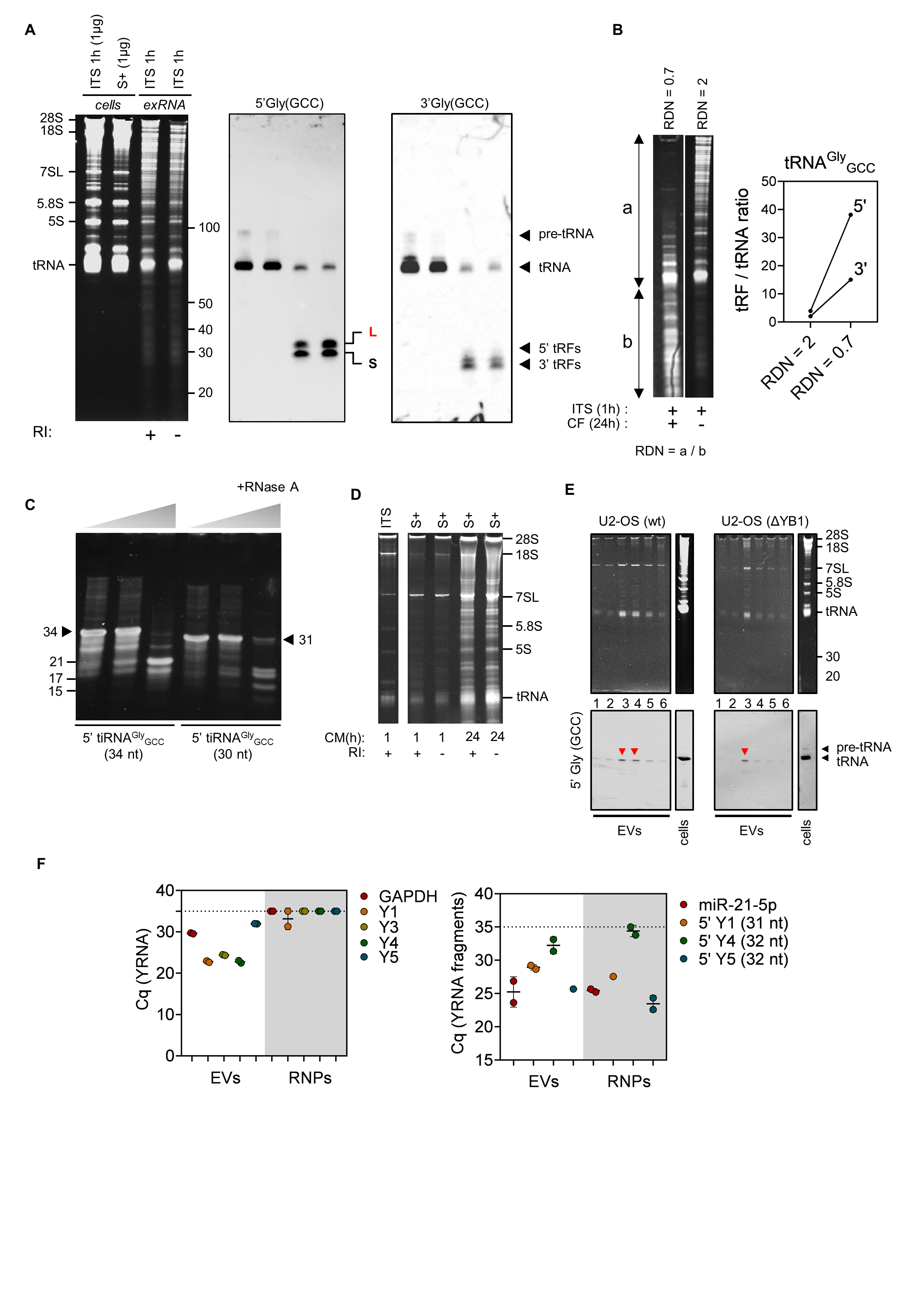

### Supplementary Fig. 5

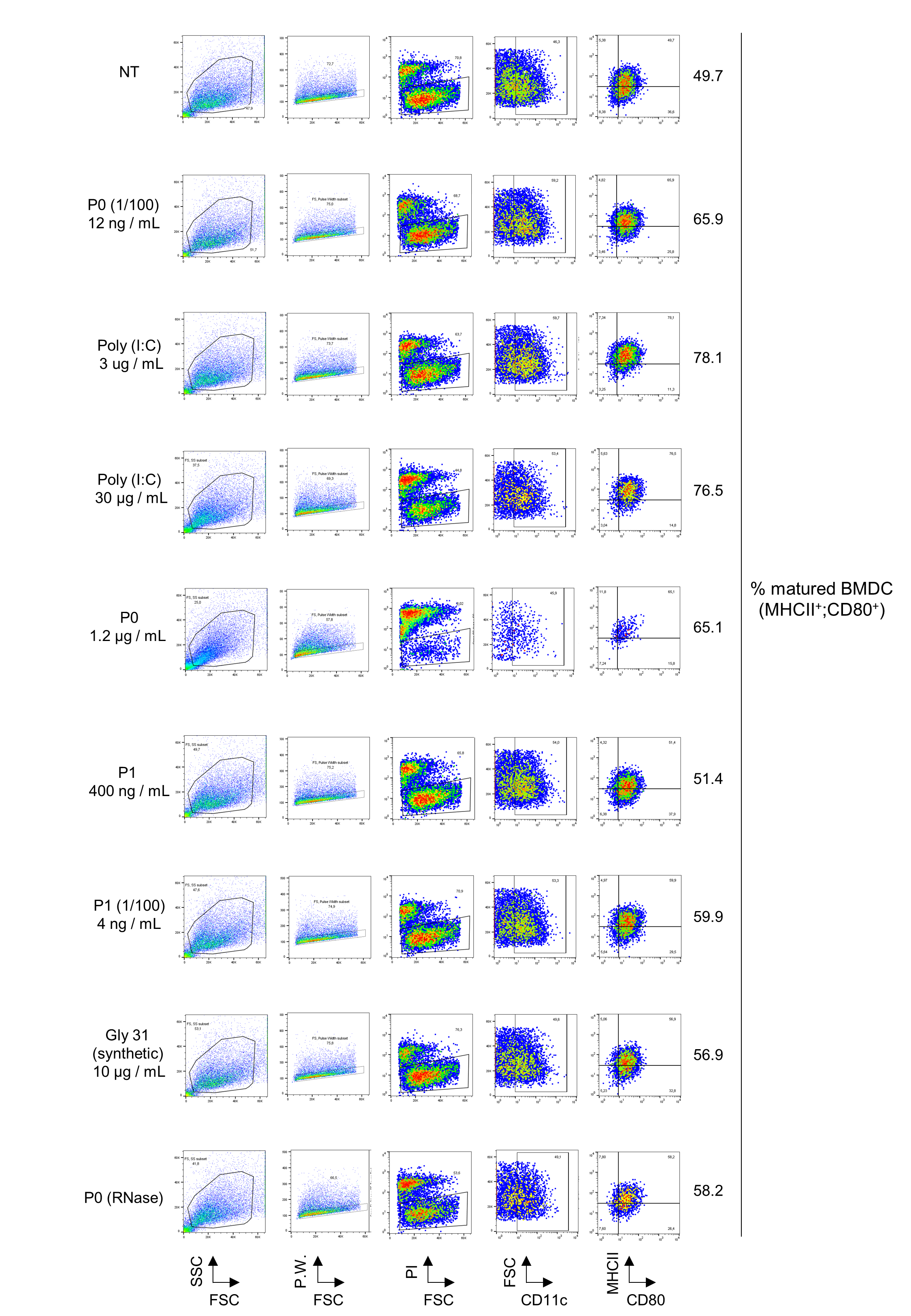
